# Modeling Protein Sequence Evolution as an Ornstein-Uhlenbeck Process in a Latent Space

**DOI:** 10.64898/2026.09.16.751972

**Authors:** Matteo De Leonardis, Andrea Pagnani

## Abstract

High-throughput directed evolution produces longitudinal sequence libraries that are ideal for probing local fitness neighborhoods but often underpowered for global inference tasks such as contact prediction. We present an unsupervised inference model that integrates directed-evolution sequencing time series with natural homologs. We project sequences into a low-dimensional latent space [1] learned from the natural multiple sequence alignment and model the experimental process as an Ornstein–Uhlenbeck dynamics in that space. Maximum-likelihood estimation of the latent drift and noise parameters determines a stationary Gaussian distribution, which induces an effective Potts model in sequence space. The inferred couplings improve structural contact prediction by combining global evolutionary constraints from nature with local, experiment-specific signals. Experiments on PSE1 *β*-lactamase and dihydrofolate reductase demonstrate the ability to identify correct complementary contacts not recovered by methods using either natural or experimental data alone, with gains concentrated in intermediate- and long-range contacts.

## 1 Introduction

High-throughput sequencing techniques paved the way for powerful biotechnological tools to study genotype-phenotype mapping for properties of interest in biological sequences [2]. In particular, Directed Evolution (DE) experiments [3] try to reproduce *in-vitro* the evolutionary process that biological sequences undergo in nature inside living organisms. They combine random mutagenesis techniques like error-prone PCR [4] and selection strategy to explore the sequence space and test the effect of the various mutations on the sequence’s fitness. DE experiments are a fundamental tool to explore the evolutionary trajectories of evolving sequences [5–7] and also to design optimal sequences to carry out specific functions [8, 9]. Despite the great benefits that this tool can provide, it still lacks a unified and general inference method to analyze its data. The most common approach consists of employing a well-established inference method for homologous proteins and applying it to the last experimental round [10, 11]. Other approaches try, instead, to describe the evolutionary process as specifically designed Markov Chain models [12–14] or standard population dynamics models [15, 16] and use the data to fit the evolutionary dynamics of the experiment. In essence, the abbreviated time scale and limited diversity of Directed Evolution experiments make them exceptionally efficient for locally mapping the fitness landscape around a starting sequence. However, this very local focus is also a limitation: any structural prediction method based solely on this experimental data achieves significantly worse performance than approaches that integrate broader homology information from natural sequences.

To bridge this gap, we introduce a method that synergistically combines DE data with information from natural homologs. Our approach involves projecting both sequence types into a common, low-dimensional latent space. This projection captures fundamental fitness constraints, allowing us to relate the diverse natural sequences to the more localized experimental variants. Within this latent space, we model evolution as an Ornstein-Uhlenbeck (OU) process [17, 18]. This dynamical model is adept at describing a system undergoing random fluctuations while being pulled toward an optimal state—perfectly capturing the balance between random mutation and directional selection in a DE experiment [19]. The process is formally described by the following stochastic differential equation:

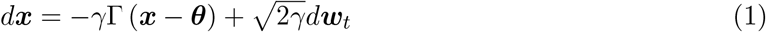

where the position of the particle is represented by the vector ***x*** ∈ ℝ^*d*^ (*d* is the latent space dimension), the matrix Γ ∈ ℝ^*d×d*^ is the external drift, ***θ*** ∈ ℝ^*d*^ is the equilibrium point, *γ* ∈ ℝ is the is the inverse characteristic time-scale of the process and ***w***_*t*_ ∈ ℝ^*D*^ represents the standard Weiner process.

The OU process provides a minimal continuous-time description of laboratory evolution in the latent space. Random mutagenesis induces diffusive exploration, while selection generates a systematic drift toward high-fitness regions. In the latent representation defined by natural sequences, this interplay corresponds to a mean-reverting stochastic dynamics. Crucially, the process admits a stationary Gaussian distribution, which we interpret as the global fitness constraints encoded by the family of natural homologs. In this way, short-timescale experimental trajectories are modeled as transient relaxations toward a natural evolutionary manifold, linking local laboratory exploration to long-term evolutionary structure.

We first define a low-dimensional (viz., latent) representation of sequence space using principal components derived from a multiple sequence alignment (MSA) of natural homologs [20] as described in Sec. 4.1. This latent space captures the dominant evolutionary constraints encoded in natural variation. Experimental sequences, including the wild type **x**_0_, are projected onto these coordinates. We then model the laboratory evolution process directly in this latent space as OU dynamics. Although individual trajectories are not observable, the population snapshots collected at each round define a time series whose likelihood can be computed analytically from the OU transition kernel *P* (**x** | **x**_0_, *t*) [21]. Maximizing this likelihood yields the parameters of the latent dynamics and their stationary Gaussian distribution. Projecting this equilibrium distribution back to sequence space induces an effective Potts model, whose inferred couplings *J*_*ij*_(*a, b*) are subsequently used for structural contact prediction within the standard Direct Coupling Analysis framework [22–25].

To assess the predictive performance of our framework, we analyze two directed evolution datasets: *β*-lactamase PSE1 [11] and dihydrofolate reductase (DHFR) [12]. In both cases, variants were selected for antibiotic resistance in *E. coli* (ampicillin for PSE1 and trimethoprim for DHFR). Although the experimental protocols are similar, the resulting sequence libraries differ markedly in their diversity. PSE1 accumulates on average 1.7 mutations (Hamming distance from the wild type) per round, whereas DHFR accumulates only 0.79. Both experiments explore a substantially narrower region of sequence space than natural homologs, yet the higher mutational rate in PSE1 generates a stronger statistical signal for structural inference. These two datasets provide complementary test cases, testing the predictive capability of our model in regimes of relatively high and low effective mutation rates.

Tab. 1**(a)** and **(b)** summarize key statistics of natural and experimental sequence datasets for DHFR and PSE1. Here, we disregard the temporal structure of the DE experiments and aggregate all variants generated within each experiment.

**Table 1.**
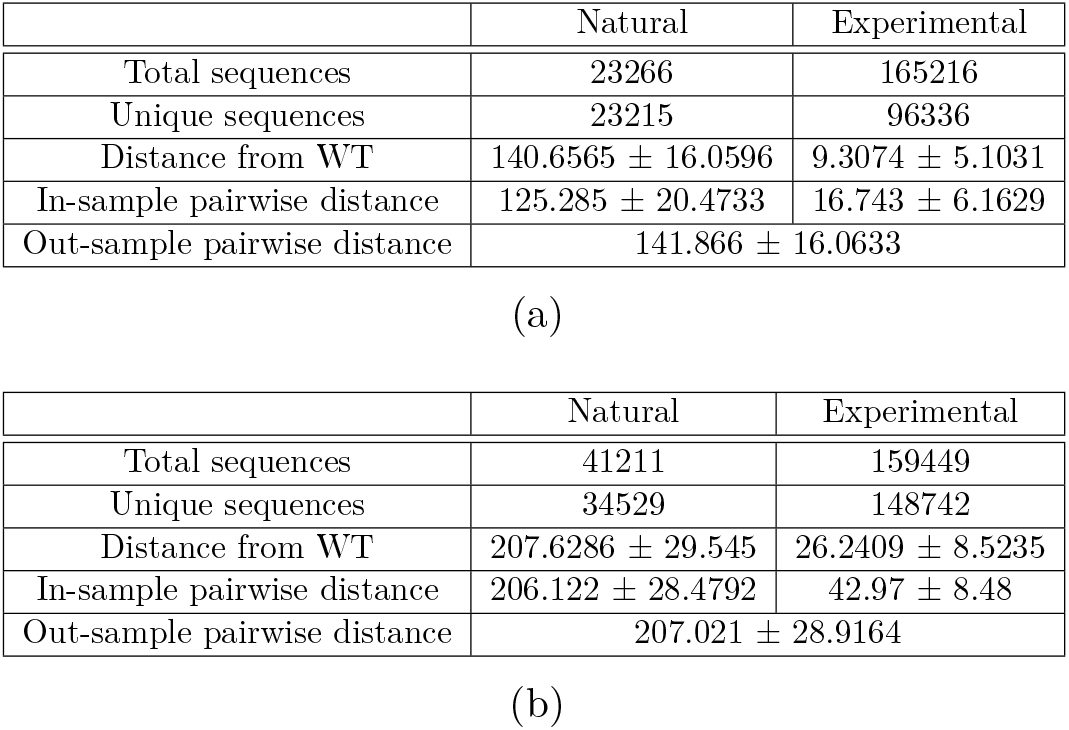
Summary statistics of natural and experimental sequence datasets for (a) DHFR and (b) PSE1. Reported quantities include the total and unique number of sequences, the average Hamming distance from the wild type (WT), the average in-sample pairwise distance, and the average pairwise distance between natural and experimental sequences (out-sample). Distances are reported as mean ± standard deviation. Experimental libraries are substantially larger than natural datasets but display markedly reduced diversity, as reflected by their small distance from WT and low internal pairwise distances. In contrast, the distance between experimental and natural sequences is comparable to the natural pairwise distance, indicating that experimental variants, while close to WT, remain far from the bulk of natural homologs.

High-throughput sequencing enables the exploration of sequence libraries that are orders of magnitude larger than the set of available natural homologs. However, despite their size, experimental libraries exhibit substantially lower sequence diversity, which may limit their usefulness for inferring global properties of the sequence landscape.

In both systems, the average Hamming distance between the wild type (WT) and experimental variants is dramatically smaller than the typical pairwise distance between natural homologs. This indicates that experimental sequences are concentrated in a narrow region of sequence space surrounding the WT.

At the same time, the average distance between experimental and natural sequences is large and comparable to the typical pairwise distance within the natural ensemble. This shows that, although experimental variants remain close to the WT, they are generally far from most natural homologs. Taken together, these observations suggest that experimental libraries probe a localized, WT-centered region of sequence space that only negligibly overlaps the broader and more diverse distribution spanned by naturally evolved sequences.

## 2 Results

We begin by projecting all experimental sequences into the latent space defined by the principal components of natural homologs (see sec. 4.3 for details). This mapping assigns to each sequence a linear projection in the *d*-dimensional latent space, allowing us to track how sequence populations evolve across experimental rounds.

As mutations accumulate, variants diffuse through this latent space, generating trajectories that reflect the underlying evolutionary dynamics. In this Principal Component representation shown in Fig. 1, we can observe how for both mDHFR, and PSE1 experiments (panel **(a)**, and **(b)** respectively), the population evolves from an initial point (the red dot representing the wild-type sequence) and progressively spreads and drifts as diversity increases during the experimental rounds of *in vitro* evolution.

**Figure 1.**
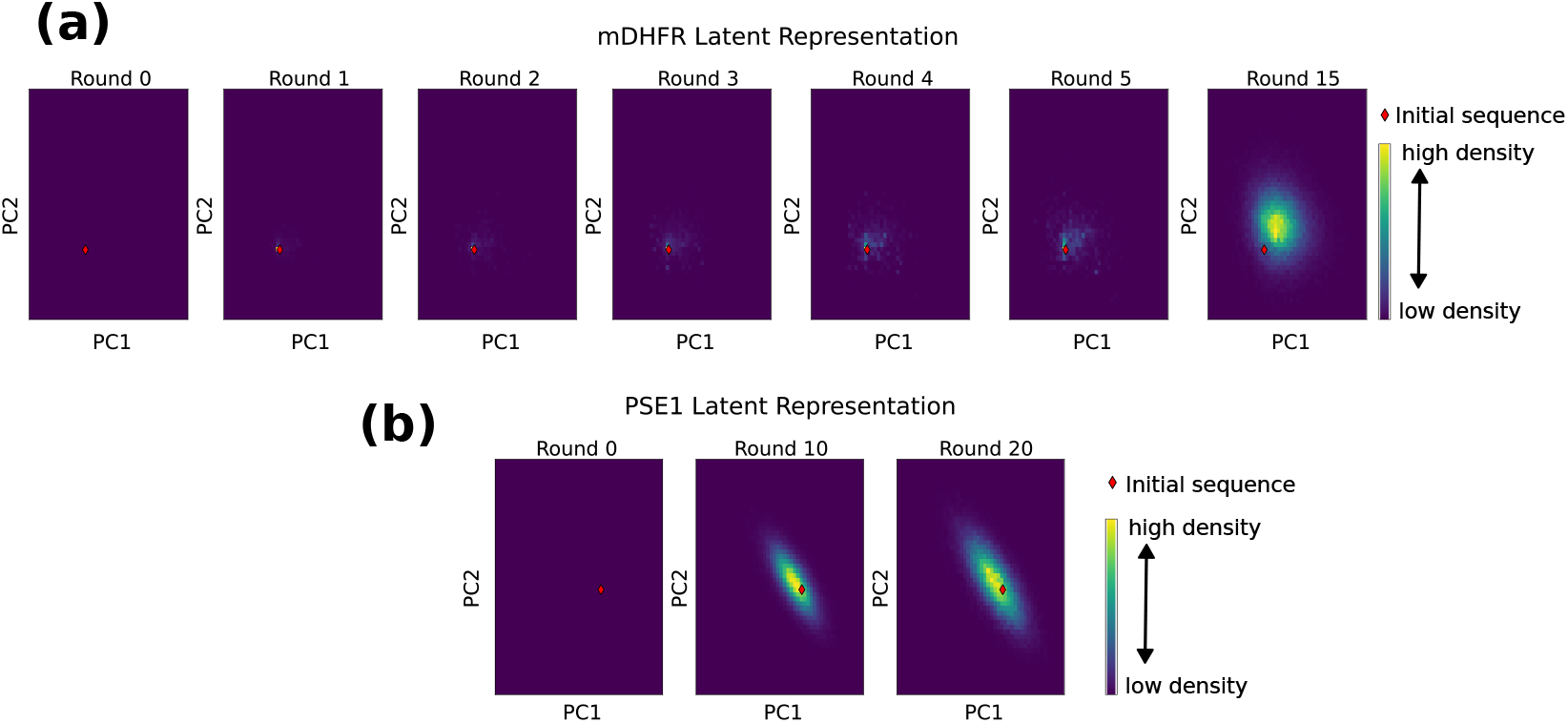
Time-series of laboratory evolution trajectories projected onto the latent space defined by the first two principal components (PC1 and PC2) derived from natural homologs. Each point represents a sequence, the color scale indicates the local density of variants in the latent space (from low to high), and the red dot is the projection onto PC1 and PC2 of the wild-type sequence. Panel **(a)** shows experimental data for mDHFR. Panel **(b)** shows for PSE1. At Round 0, only the wild-type sequence is present, resulting in a single point (red diamond, labeled *Initial sequence*). Over successive rounds, the distribution progressively diffuses across the latent space.

This observation is consistent with our modeling assumptions. The OU process describes diffusion in the presence of a single attracting point, leading to unimodal, mean-reverting dynamics in latent space.

While principal component analysis applied to heterogeneous datasets can produce multiple, well-separated clusters, in our case, the latent space is defined using natural homologs, which exhibit substantially greater diversity than the experimentally generated variants. As a result, all experimental sequences remain confined to a single region of the latent space, forming a compact, unimodal distribution centered around the wild type (see fig. 6). This property is essential for the applicability of the OU model, which cannot capture multimodal evolutionary dynamics under Eq. (1).

### 2.1 Predicting Contacts

We use the time-series data in this latent space to infer the external force Γ and the equilibrium point ***θ*** using Maximum Likelihood, as described in Sec. 4.5, and then use the mapping from sequence space to latent space to infer a set of Potts model parameters, as customary in DCA-like approaches, that assign the probability

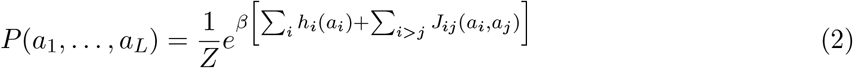

to each sequence ***a*** = (*a*_1_, …, *a*_*L*_) of length *L* and *Z* is the normalization factor.

PlmDCA [26] is an inference method to estimate Potts parameters from a set of naturally evolved sequences, based on a pseudo-likelihood approximation. It is considered a standard approach for this task; when many sequences are available, it achieves contact prediction performance comparable to the Boltzmann learning approach. We train PlmDCA on natural sequences to obtain reliable structural contact predictions and use these predictions to quantify the additional information that our OU model can infer compared to a DCA-like model that does not use experimental sequences. Since the diversity of natural sequences is much greater than that of experimental sequences, DCA is expected to yeld more accurate predictions than any inference method trained only on experimental data. In the Supplementary Information, Fig. S1 shows that PlmDCA achieves very accurate predictions on both datasets of natural sequences. Here, we are instead interested in assessing whether training our model on experimental data provides information not already included in PlmDCA and, in some way, complements contact prediction by leveraging patterns observed in laboratory-designed variants.

Our model, based on OU dynamics, leverages time-series data in the latent space to estimate the parameters Γ and ***θ*** defined in Eq. 1. At equilibrium, the mapping between sequence space and the latent representation allows one to relate Γ to the coupling matrix 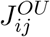(*a, b*) that encodes pairwise interactions in the Potts model, as commonly used in statistical modeling of protein sequences. Here, *J*_*ij*_(*a, b*) is a four-index object with site indices *i, j* = 1, …, *L* and amino-acid states *a, b* = 1, …, *q*, which can be equivalently represented as a matrix of size *Lq* × *Lq* by combining site and state indices. The mathematical details of this procedure are provided in Sec. 4.3. The resulting expression for the couplings is given by:

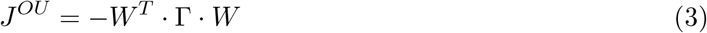

where *W* is the PCA projection matrix, of size *d* × *Lq*, which maps sequences onto the *d* principal components computed from natural homologs.

Looking at how we derive the coupling matrix of the inferred Potts model, one can notice the analogy with another inference method known as the *Gaussian Approximation* in the context of DCA [27]. In fact, the Gaussian Approximation estimates the coupling matrix as the inverse of the empirical covariance matrix *C* [28]:

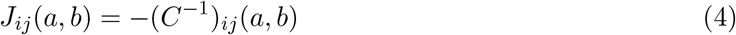

Eq. (3) expresses how we estimate the Potts model coupling matrix *J* from the external force Γ and coincides with a low-rank expression of the covariance matrix computed on the data if Γ is a diagonal matrix whose elements are the inverse of the *d* largest eigenvalues of the covariance matrix:

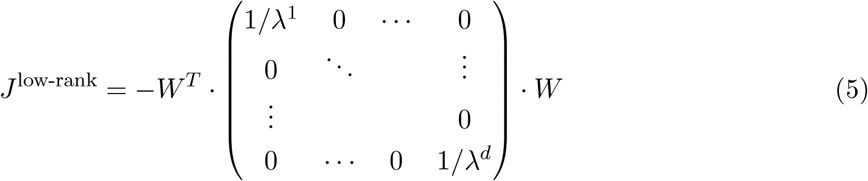

Instead, our OU-based model can infer a generic (positive-definite) matrix that also weights the covariance of different PCA patterns according to the time-series data of the experiment. Therefore, it appears natural to validate our approach by comparing the contact predictions from our model with the low-rank representation of the Gaussian Approximation for DCA. We stress that our aim is not to provide a better alternative that outperforms standard DCA approaches, because our method performs an essentially different task. DCA fits the statistics of naturally evolved sequences to identify relevant residue–residue interactions that may signal contacts in the 3D structure. Our aim, instead, is to enrich the information obtained from natural sequences and, thanks to experimental evolution data, detect additional contacts not found by standard DCA approaches.

Fig. 2 **(a)** and **(b)** show how the contact predictions from our model compare with the low-rank Gaussian approximation mentioned above, for mDHFR and PSE1, respectively. Throughout this paper, we adopt the customary threshold of *L/*2 top-ranked pairs as predicted contacts. A threshold of *L* is also common in these methods; the exact choice is somewhat arbitrary, and no universally accepted standard exists.

**Figure 2.**
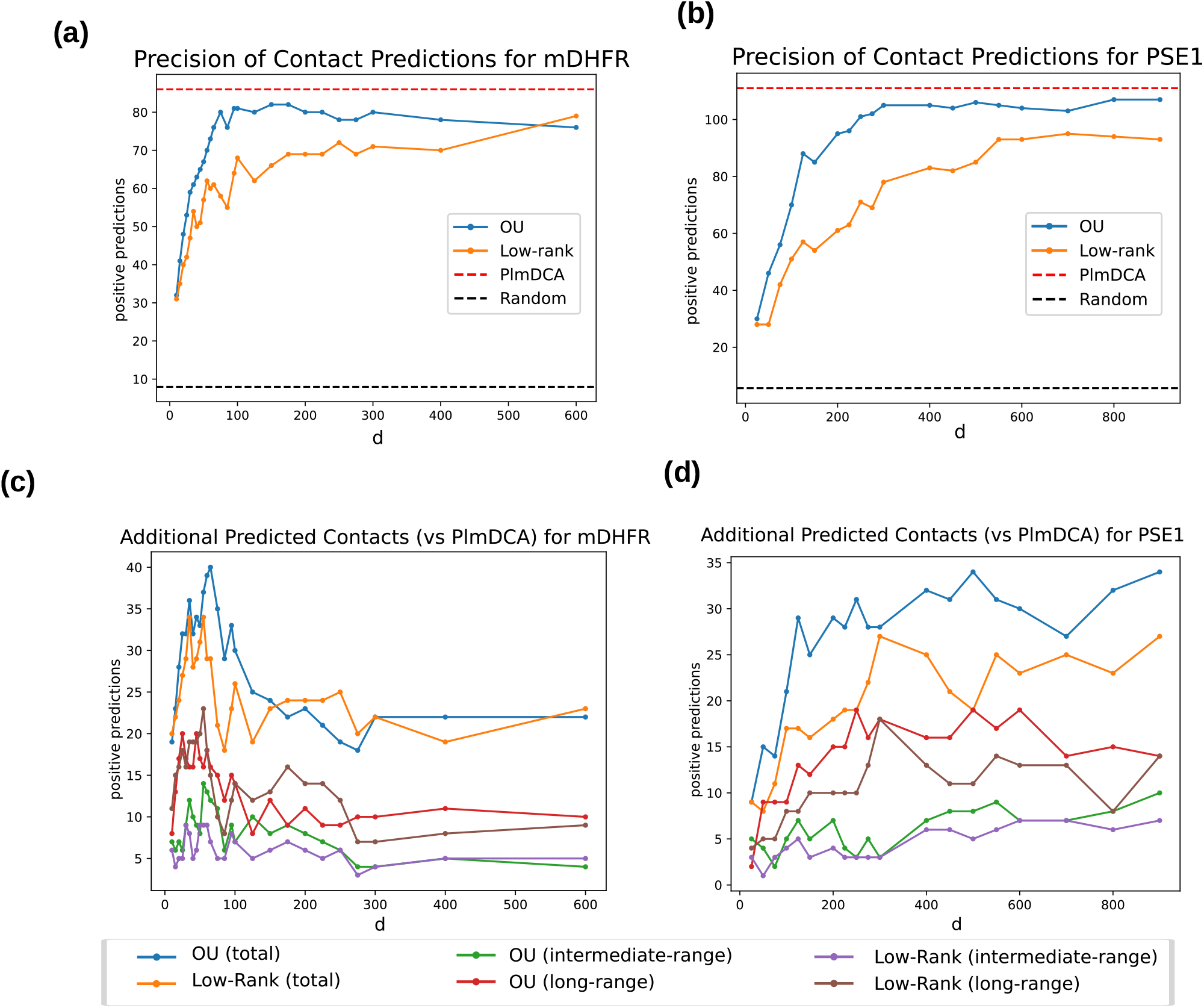
Panel **(a)**: On the y-axis, we display the number of positive predicted contacts among the *L/*2 best-ranked pairs for our OU model (blue curve) trained on mDHFR experimental data and the respective low-rank Gaussian approximation (orange curve). The x-axis value *d* is the dimension of the latent space representation, for the OU model, and the maximum rank of the precision matrix computed in the Gaussian approximation. The red dotted line indicates the number of correct contact predictions from PlmDCA trained on natural mDHFR sequences. Panel **(b)**: same as panel **(a)** but for PSE1. Panel **(c)**: The y-axis indicates the number of additional correct predictions with respect to PlmDCA trained on natural mDHFR variant as a function of the latent space dimension *d*; in the OU model case, they represent the dimensionality of the latent space, while in the low-rank Gaussian approximation, they represent the maximum rank of the precision matrix computed from the data. The blue line refers to additional contacts found by the OU model, the orange line to the low-rank Gaussian approximation. The other lines indicate, according to the legend, how many of these additional contacts are found to be at long-range (|*i* −*j* |*>* 23) or at intermediate-range (12 *<* |*i*− *j*| ≤23) since they are considered to be more structurally significant. Panel **(d)**: same as Panel **(c)** but for PSE1.

The blue curve indicates the number of correct contact predictions from our OU-based model, while the orange one refers to the low-rank Gaussian approximation. The red dashed line indicates the performance of PlmDCA. The x-axis reports the values of the parameter *d*, which in our case is the dimension of the latent space in which experimental sequences evolve, while in the Gaussian approximation it represents the rank of the inverse covariance used to infer *J* ^low-rank^ in Eq. (5).

The figures show that, in both cases, restricting the number of PCA components considered in the covariance matrix reduces the number of predicted contacts with respect to PlmDCA. For both approaches, increasing *d* significantly improves contact prediction accuracy. In particular, for mDHFR we recover performance comparable to PlmDCA for *d* ≈ 150, which corresponds to approximately 55% of the explained variance from the PCA components; for PSE1, this occurs at *d* ≈ 300, corresponding to approximately 65% of the total variance of natural sequences.

Moreover, the figures show that the number of correctly predicted contacts from the OU model is consistently higher than that of the low-rank Gaussian approximation. This means that while restricting the number of PCA components typically decreases contact prediction accuracy, incorporating the patterns of experimental sequences can preserve information from relevant PCA components and recover part of the predictive power. This trend is consistent for both experiments and for almost all values of *d*, with the only exception being *d* = 600 for mDHFR, where the low-rank approximation slightly outperforms the OU model.

The picture becomes more complex when we analyze only the additional contact predictions, i.e., those not already identified by PlmDCA. This analysis aims to assess whether these additional contacts arise mainly from restricting PCA components or from information related to the evolutionary dynamics of the experiment.

From fig. 2 **(c)**, we observe that the novel predictions from our OU model exceed those from the low-rank approximation up to *d* = 150. Beyond this point, the trend reverses and the low-rank approximation finds more additional contacts than the OU model. At *d* = 300, these novel predictions become almost constant and similar in value for both methods. Regarding structural significance, the number of long-range correct predictions is slightly higher for the low-rank approximation for almost all values of *d* up to 300, while the opposite holds for intermediate-range predictions.

Fig. 2 **(d)** shows the same analysis for PSE1. In this case, the picture is clearer: our OU model exhibits a consistently better predictive capacity for new contacts, both long- and intermediate-range. This difference likely arises because the experiment on mDHFR is much less informative than that on PSE1. In mDHFR, the mutation rate is too low to generate sufficient diversity to produce a significant statistical signal for contact prediction. This is consistent with the findings of the reference study [12].

Although there is no well-established inference method specifically designed for laboratory evolution experiments, common practices exist for applying DCA-like methods to such data. The simplest and most standard approach consists of combining natural and experimental sequences and training PlmDCA; this is the basic idea behind the *EVCouplings* method [29]. This approach often provides meaningful predictions because sequences are reweighted according to their similarity, leading to a strong down-weighting of experimental sequences, since they derive from the same wild-type sequence. As a result, adding experimental sequences produces only minimal changes in the inferred fitness landscape: most information remains consistent with patterns from natural sequences, with only limited additional contributions from the experimental variants. In the following, we will use the term Combined-MSA (or simply Combined) to refer to this approach, which follows the same general strategy as EVCouplings: experimental sequences are appended to the MSA of natural homologs. No additional alignment step is required, due to the way the MSA of natural sequences was constructed (see Sec. 4.1). PlmDCA is then applied to the resulting alignment using the default sequence reweighting.

Fig. 3 panels **(a)** and **(b)** show the comparison of the additional contacts found using Combined-MSA with respect to PlmDCA and our OU model. The additional predicted contacts are very limited, especially for mDHFR, and our OU model gives predictions that are more complementary with respect to PlmDCA. In panel **(a)** it is evident that adding experimental variants to the training set for PlmDCA is in practice irrelevant due to the very limited diversity obtained in the experiment. For PSE1, instead, see panel **(b)**, adding experimental variants changes slightly the predictions and our OU model is able to provide more diverse predictions provided that the dimensionality of the latent space is not too small. Fig. S2 shows the locations along the sequence of these additional predictions obtained by *Combined-MSA*.

**Figure 3.**
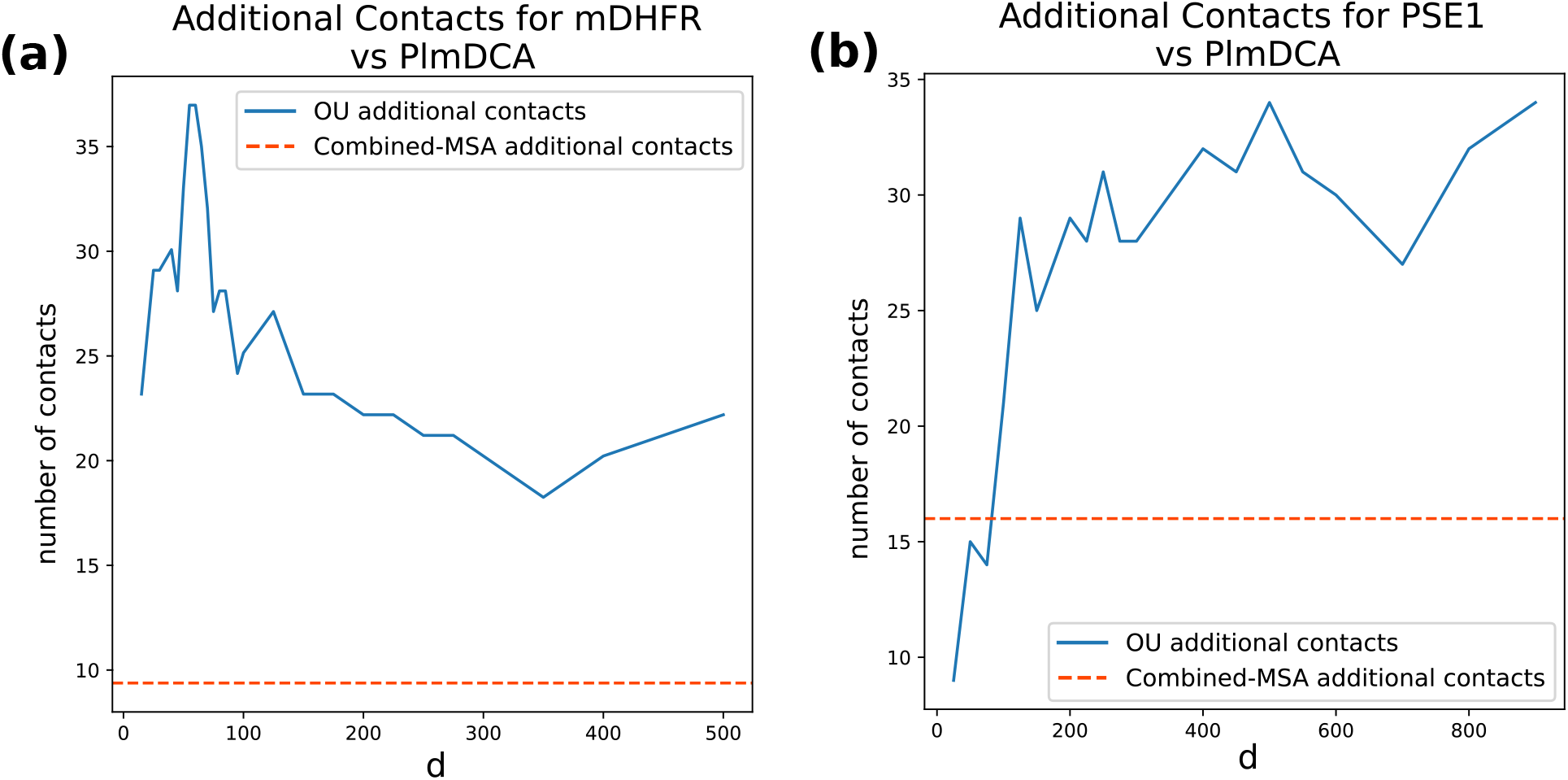
Comparison of the number of additional correct contacts predicted with respect to PlmDCA by the OU model and the *Combined-MSA* method. Panel **(a)** refers to DHFR, while panel (a) refers to PSE1. The x-axis reports the latent-space dimension *d* used by the OU model, while the y-axis reports the number of additional correct contacts with respect to PlmDCA. The orange dotted line indicates the number of additional correct contacts obtained with the *Combined-MSA* method and serves as the baseline for comparison. While *Combined-MSA* yields only a small number of additional contacts relative to PlmDCA, the OU model identifies substantially more additional contacts for almost all values of *d*, with the exception of PSE1 at very small values of *d*.

### 2.2 Choosing Latent Space Dimension

In our analysis, tuning the hyperparameter *d* remains the main open question, since there is no clear *a priori* criterion for choosing it. However, as it can be observed in Fig. 2 panels **(a)** and **(b)**, the quality of the prediction of new contacts seems to stabilize provided *d* is sufficiently large. To make this observation more quantitative, we introduce the quantity *ρ* as the relative weight the off-diagonal norm of the inferred precision matrix, defined as

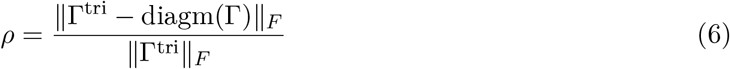

where ∥ · ∥ denotes the Frobenius norm, and diagm(Γ) is the diagonal matrix with entries equal to the diagonal elements of Γ, and Γ^tri^ is the upper triangular part of Γ. The behavior of this quantity is reported in fig. 7 for both dataset for each value of *d*.

In both experiments, *ρ* shows a similar trend: it increases with *d* and then saturates. For PSE1, saturation occurs around *d* = 500, which corresponds to a value at which the OU model already has enough principal components such that its performance is really comparable to PlmDCA and yields the maximum number of additional contact predictions, including long-range ones. For mDHFR, instead, *ρ* starts to saturate around *d* = 275. The model’s predictions become comparable to PlmDCA earlier, around *d* = 75, and the number of additional contacts reaches its maximum before saturation; however, only at *d* = 275 do the positive long-range predictions stabilize and surpass those of the low-rank approximation.

### 2.3 Predicting Evolutionary Dynamics

The contact-prediction analysis evaluates whether the stationary distribution induced by the inferred OU process contains structurally informative couplings. We next asked a complementary question: does the learned finite-time dynamics assign high probability to the experimentally observed evolutionary trajectory itself? To address this, we performed a trajectory-reconstruction analysis in which the OU transition kernel was used to score experimentally observed variants and compare them with uniformly random variants generated from the wild-type sequence (*purely mutational null model*).

We considered two versions of this analysis. A first, *full time-series analysis*, where the model was trained using the data from all the experimental rounds. This provides an in-sample control, testing whether the fitted OU dynamics is internally consistent with the observed trajectory. In the second, *held-out analysis*, the final evolutionary round was removed from the training set and used only for evaluation. This leave-last-round-out setting provides a genuine out-of-sample test of whether the inferred dynamics can recognize future evolved variants that were not used during parameter inference.

For each latent dimension *d*, we computed the OU transition score of variants in the final round and compared the score distribution of experimental sequences against that of uniformly generated random sequences. The statistical test was one-sided, with the alternative hypothesis that experimental variants receive higher scores than random variants. To generate random sequences, we use the same average mutation rate fitted from the experimental data and simulate a purely mutational process with no selection involved. Therefore, their distribution of Hamming distances from wild-type sequence match the experimental data. Small p-values indicate successful trajectory reconstruction, whereas p-values close to one indicate that the random sequences are favored by the model.

For PSE1, trajectory reconstruction is strongly supported. When the model is trained on the full time series, the direct transition from the wild type to the final round gives p-values numerically equal to zero across all tested dimensions, and the transition conditioned on the intermediate round remains highly significant, with p-values of order 10^*−*32^ − 10^*−*26^. Importantly, the same qualitative result is obtained in the *held-out* setting: the direct wild-type-to-final-round comparison again gives p-values equal to zero across all tested dimensions, and the transition from round 10 to the held-out final round gives p-values of order 10^*−*31^ − 10^*−*21^. Thus, for PSE1, the OU dynamics inferred from earlier rounds assigns substantially higher likelihood to the held-out evolved sequences than to random sequences at comparable mutational scale.

The model exhibits a milder predictive capacity on DHFR dataset. In the *full-time-series* analysis, the trajectory-reconstruction signal is strong: p-values become significant already at low dimension and are numerically zero for the direct transition from d=50 onward, with similarly small p-values for transitions conditioned on earlier rounds for *d* ≥ 40. However, because the final round is included in the training set in this analysis, this result should be interpreted as an in-sample consistency check rather than as evidence of predictive reconstruction.

The *held-out analysis* on DHFR gives a weaker result. When the final round is removed from training, most latent dimensions yield p-values close to one, indicating that random variants are often assigned higher trajectory scores than the experimental held-out variants. A significant signal appears only in a narrow range of dimensions, most notably around *d* = 125 and *d* = 150, where the direct wild-type-to-final-round p-values are 1.39 × 10^*−*55^ and 6.63 × 10^*−*28^, respectively. This suggests that, for DHFR, held-out trajectory reconstruction is sensitive to the choice of latent dimension and is not robust across the full range of PCA representations.

A visual representation of these results is shown in Fig. 4, where a threshold value of 0.05 has been chosen as significant according to p-value. A more detailed table with precise p-values is reported in Tabs. S1-S4 in Supplementary Information. The mathematical details of this analysis are described in sec. 4.7, especially for the cases reported in fig. 4 when the ancestor sequence is taken at intermediate round times.

**Figure 4.**
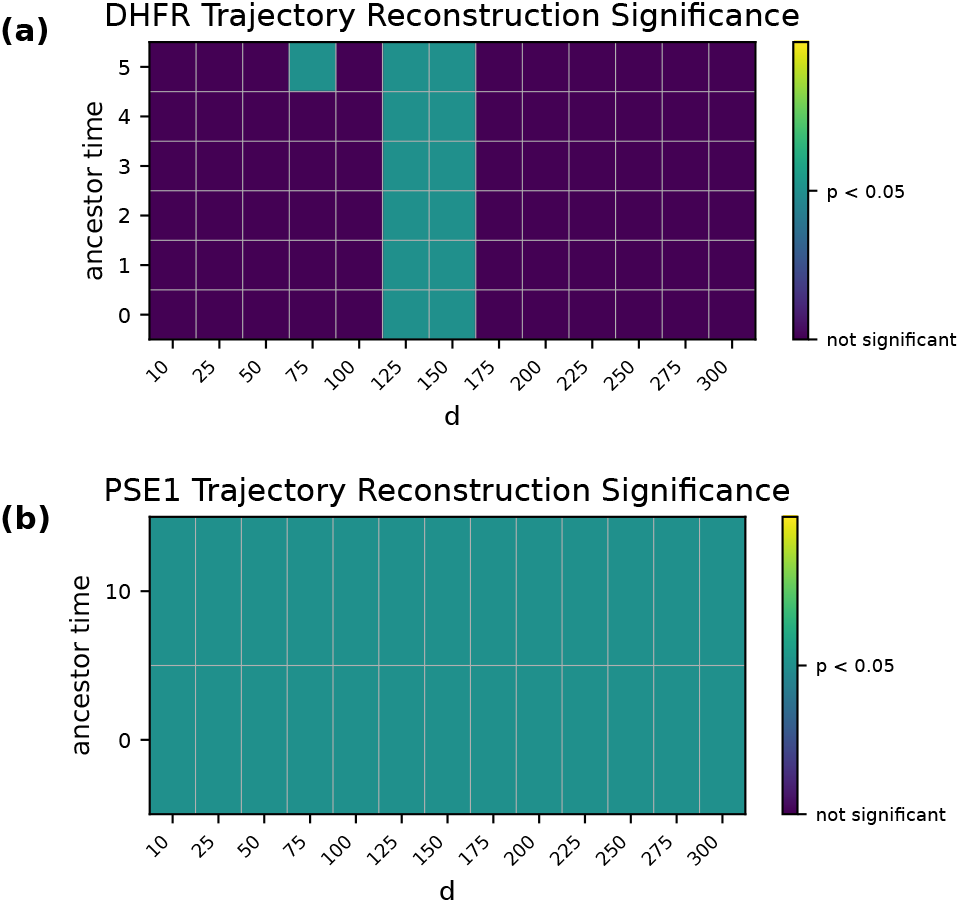
Trajectory-reconstruction significance in the leave-last-round-out setting. For each dataset, the OU model was trained after removing the final experimental round, which was then used only for evaluation. Each cell indicates whether the trajectory-reconstruction test is significant at the threshold *p <* 0.05, comparing OU transition scores of experimental final-round variants against uniformly generated random variants. The alternative hypothesis is that experimental variants receive higher transition scores. Panel **(a)** shows DHFR, where significant reconstruction is restricted to a narrow range of latent dimensions, mainly *d* = 125 and *d* = 150, with additional isolated signal at *d* = 75 *t* = 5 ancestor time. Panel **(b)** shows PSE1, where reconstruction is significant across all tested latent dimensions and ancestor times.

Overall, the trajectory-reconstruction analysis supports the dynamical interpretation of the model most clearly for PSE1. In this dataset, the final evolved population is recognized by the OU dynamics even when it is excluded from training in the *held-out analysis*.

For DHFR, the in-sample reconstruction indicates that the model can fit the observed trajectory when all rounds are available, but the *held-out analysis* reveals limited predictive power except in a narrow range of latent dimensions. This difference is consistent with the lower diversity of the DHFR experiment and with the broader conclusion that dynamical inference from directed-evolution data requires sufficient mutational spread to generate a statistically informative signal in latent space.

To assess the robustness of this result, we repeated the *held-out analysis* using more stringent null models that preserve progressively more information from the experimental sequences. In the purely mutational case (no fitness, *β* = 0 in Eq. (2)), the probability that each position is mutated is matched to the corresponding experimental round, while the identity of the mutant amino acid is randomized. Under this control, and using the wild-type as the initial sequence, significant trajectory reconstruction is obtained for PSE1 when a sufficiently large latent representation is used, approximately *d* ≥ 400, but not for DHFR. We also considered a purely mutational *site-dependent profile* model, in which each position is sampled according to its empirical amino-acid frequencies. In this case, experimental and randomized sequences are generally not significantly separated: this stricter null model reproduces much of the variation observed after projection into the latent space, indicating that a substantial fraction of the trajectory-reconstruction signal is encoded in the single-site sequence statistics. Complete results for these null models are reported in the Supplementary Information.

## 3 Conclusion

In this work, we introduced a unified framework to integrate Directed Evolution (DE) data with information from natural sequences by modeling evolutionary trajectories in a latent space defined by PCA through an OU process. This construction provides a principled way to connect the local exploration of sequence space performed in laboratory experiments with the global constraints encoded in naturally evolved families. By mapping the inferred stationary distribution back to sequence space, we obtained an effective Potts model whose couplings can be directly compared with standard DCA approaches.

Our results highlight a clear contrast between the two experimental systems analyzed. In the case of PSE1, the DE experiment generates sufficient sequence diversity to produce a measurable statistical signal in the latent space. In this regime, the OU-based inference consistently improves over a simple low-rank Gaussian approximation and yields additional structurally meaningful contacts that are not already captured by PlmDCA. The improvement is particularly evident for long- and intermediate-range contacts, indicating that the temporal information encoded in the experimental rounds provides complementary constraints beyond those present in natural sequence variation alone. In this setting, the dynamical modeling of the experiment effectively extracts information that enriches that derived from natural sequences and would otherwise remain hidden. In contrast, the experiment performed on mDHFR operates in a markedly lower-diversity regime.

The limited mutational spread around the wild-type sequence results in a weak statistical signal in the latent space, which in turn constrains the amount of information that can be reliably extracted by our dynamical model. Although The OU approach performs at least as well as the corresponding low-rank approximation in terms of total correct contactsfor nearly all tested dimensions, the number of additional structurally significant predictions remains limited and does not show a robust advantage, also for-long range interactions. Moreover, the novel contacts inferred for mDHFR display a consistent enrichment compared to low-rank Gaussian approximation only for small values of *d* when experimental sequences are related to a few limited sequence patterns. Overall, the experimental data for mDHFR do not provide a statistically strong enough perturbation of the natural covariance structure to enable confident, additional structural predictions.

The effectiveness of dynamical inference from DE data critically depends on the diversity generated during the experiment. When the mutational rate is sufficiently high, as in PSE1, the OU framework can leverage the time-series information to extract complementary structural constraints. When diversity is too limited, as in mDHFR, the experimental data do not carry enough independent statistical signal to significantly refine or extend predictions derived from natural sequences alone. Using PCA main directions from natural sequences as coordinates of the latent space has indeed several advantages. First, it can be easily and efficiently computed from the data, and, in addition, experimental sequences cover just a tiny portion of the PCA space outlined by natural sequences. This avoids the formation of multiple clusters along evolutionary trajectories that couldn’t be described by OU dynamics. Nevertheless, this choice comes with some disadvantages. The main one is that the order of the principal components from PCA is completely unrelated to the experimental variants in a specific experiment. Adding more PCA directions can be completely uninformative and unrelated to the experiment, and makes it harder for the model to extrapolate the dynamical patterns driven by selective pressure. This seems not to be an issue for the experiment on PSE1 and, reasonably, for those experiments that explore the sequence space in proximity of the wild-type. On the other hand, the experiment performed on mDHFR shows this issue. The predictive quality increases and decreases according to the PCA coordinates that we add to the latent space representation, producing local optima that depends on the specific patterns we are adding to the dynamics. Future work should indeed address this issue and improve how we construct the latent space using only those sequence patterns that are related to the experimental variants.

In addition to contact prediction, the trajectory-reconstruction analysis provides a direct validation of the dynamical component of the model. By comparing the OU transition scores of experimentally evolved variants with those of others that were randomly generated from a purely mutational process, we tested whether the inferred dynamics assigns higher likelihood to sequences that actually appear along the evolutionary trajectory. The results are strongest for PSE1, where the held-out final round is consistently distinguished from random sequences, supporting the idea that the OU process captures a genuine dynamical signal in the latent space. For mDHFR, the signal is weaker and strongly dependent on the latent-space dimension, reinforcing the conclusion that limited mutational diversity constrains the amount of information that can be extracted from the experimental time series. Overall, this analysis complements the structural-contact results by showing that, when sufficient diversity is present, the model not only improves inference of sequence constraints but also captures predictive information about the temporal evolution of the tested protein sequences.

## 4 Methods

### 4.1 Constructing MSA from natural homolgs

As mentioned in the Introduction, this method exploits natural sequences to define the coordinates of the latent space onto which the experimental sequences are projected. To ensure consistency, natural sequences must be aligned to the experimental wild-type (WT) sequence (and, consequently, to its variants).

We used jackHMMER [30] to retrieve homologous sequences from the PF13354 and PF00186 families for *β*-lactamase and DHFR, respectively, using the corresponding experimental WT sequences as queries. From the resulting MSA, we retained only those columns corresponding to match states in the alignment of the experimental WT sequence. This procedure ensures that only positions aligned to residues of the experimental WT are preserved. We then filtered out sequences containing more than 50% gaps or invalid characters.

The resulting MSA has the same length as the experimental WT and contains, at each position, either an amino acid if the residue in the natural sequence aligns to the corresponding WT position, or a gap otherwise.

### 4.2 Latent space mapping

We performed principal component analysis (PCA) on the produced MSA of natural homologs to extract the dominant modes of natural sequence variation (see fig. 5). The matrix *W* is defined as the linear projector onto the first *d* principal components.

**Figure 5.**
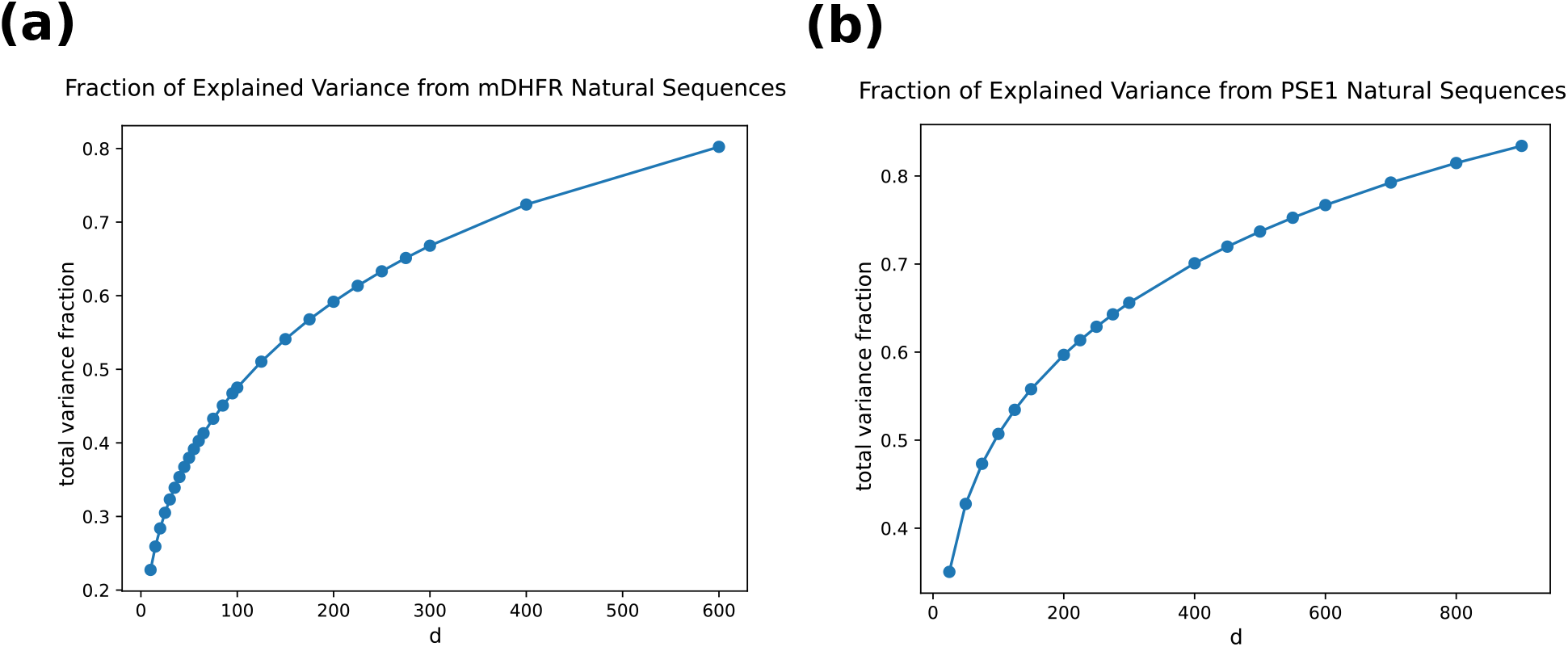
Percentage of variance explained by the first *d* components of PCA computed on natural sequences. (a) for mDHFR and (b) for PSE1 natural sequences.

We do not apply sequence reweighting when computing PCA on the natural MSA. While reweighting is standard in covariance-based methods to correct for phylogenetic biases, in our framework PCA is not used to estimate statistical couplings but only to define a linear embedding of sequences into a latent space. The subsequent OU inference step assigns different weights to latent directions through the matrix Γ, effectively selecting the relevant modes for the experimental dynamics. Consistently with this interpretation, we find that the precise choice of PCA components has only a limited impact on the results. In particular, we repeated the same analysis selecting *d/*2 components associated with the largest eigenvalues and *d/*2 components associated with the smallest ones [31], obtaining essentially the same performance. This suggests that the detailed structure of the PCA basis is not a critical factor for the modeling, although improving the construction of the latent representation remains an interesting direction for future work.

**Figure 6.**
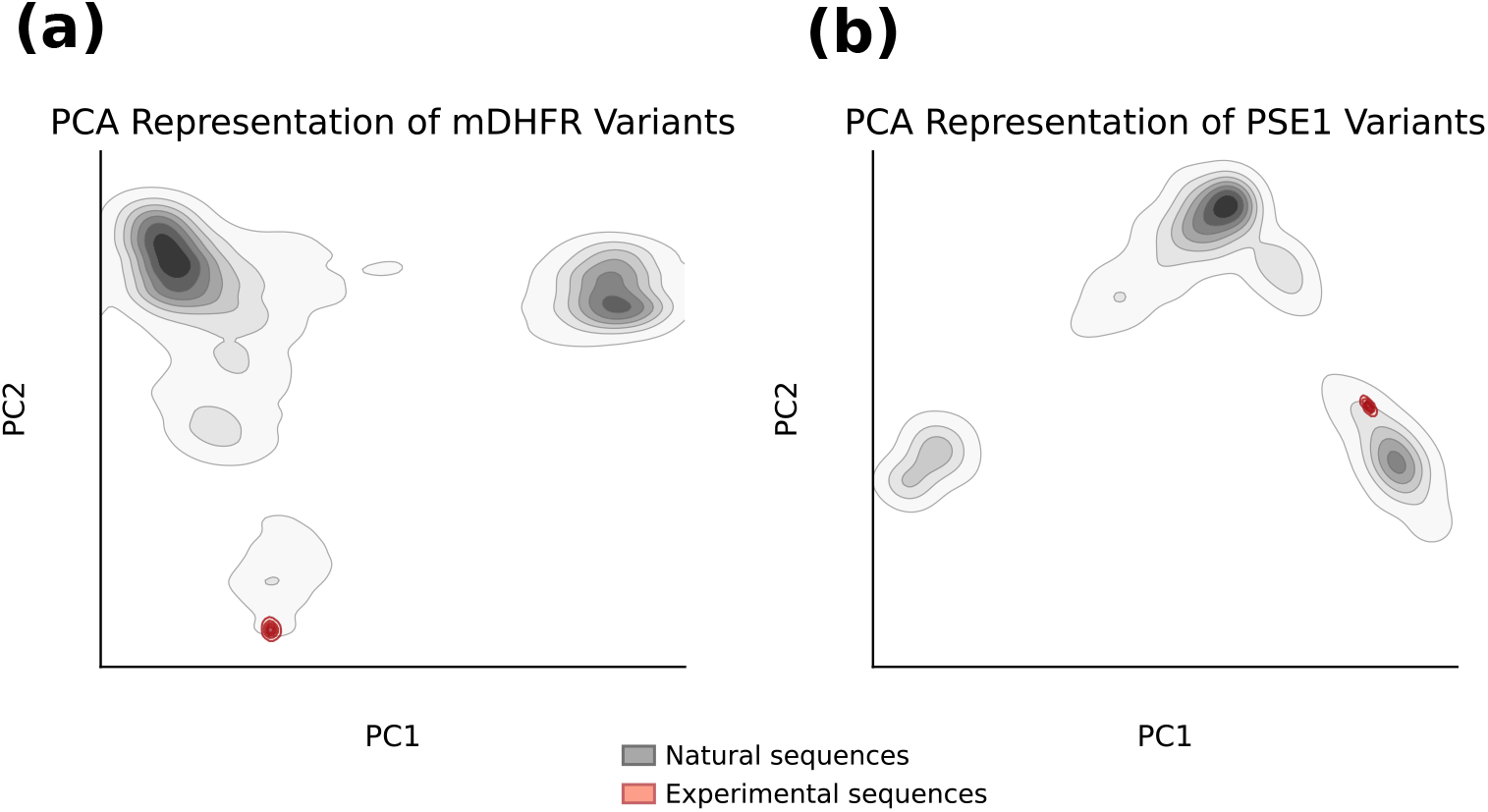
Two-dimensional PCA projections of sequence variants for **(a)** mDHFR and **(b)** PSE1. Gray contours represent the density of natural sequences derived from multiple sequence alignments, while red contours indicate the distribution of experimentally tested variants. In both cases, experimental sequences occupy a restricted region of the broader natural sequence landscape, highlighting the limited portion of sequence space explored by directed evolution experiments.

### 4.3 Projecting sequences into low-dimensional space

Protein sequences are assumed to be part of a multiple sequence alignment of homologous proteins. For this reason, we can consider a protein sequence of length *L* represented as **a** = (*a*_1_, …, *a*_*L*_), where at each alignment position, *a*_*i*_ takes values in an alphabet of size *q* = 21 (20 amino acids plus the gap symbol). We use a one-hot representation arranged into *L* consecutive blocks of length *q*, one block per site. For site *i*, the component corresponding to symbol *a* is defined as

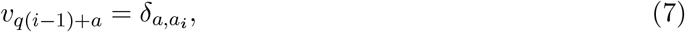

so that exactly one entry in each block equals 1 and all the others are zero.

We introduce a linear mapping *W* ∈ ℝ^*d×Lq*^ from the space of one-hot encoded sequences ℝ^*Lq*^ to a *d*-dimensional latent space ℝ^*d*^. Given a sequence represented by its one-hot vector ***v***, its latent representation is defined as ***x*** = *W* ***v***. The rationale behind this construction is that the latent space captures the relevant collective degrees of freedom governing sequence variability. We assume that the distribution of latent variables follows a Boltzmann law. In particular, we consider a quadratic energy function, so that the corresponding equilibrium distribution in latent space is a multivariate Gaussian.

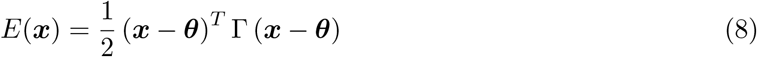

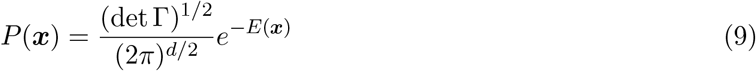

We can use the mapping *W* to backtrack each point in the latent space to the *Lq*-dimensional sequence space and define an energy function in the latter according to the transformation ***W*** and eq. 8. We obtain the following expression of the energy function in the sequence space

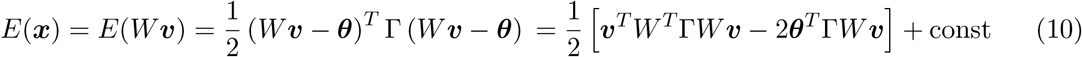

This energy function defines a Potts model with pairwise interactions in the sequence space [32], where the external fields are given by

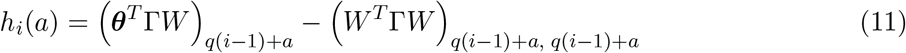

and the coupling matrix assumes the following expression (already introduced in eq. 3)

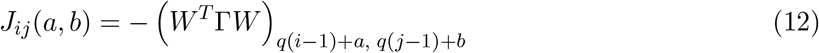

that consists in a low-rank representation that assigns the probability of each sequence based only on the latent space representation.

Now, strictly speaking, the energy function derived in eq. 10 describes an equilibrium distribution over the space ℝ^*Lq*^, but we are interested only in its restriction to one-hot encoded vectors. We do not address this issue directly, as we are not aiming to sample sequences using this Potts model. Instead, we rely on data to predominantly weight meaningful sequences. Therefore, we interpret the parameters {***h*** *J*} as the fields and couplings of a Potts model describing the equilibrium of an interacting system of Potts variables. One evident issue with this interpretation is that there is no way to avoid self-couplings (diagonal terms in the *J*_*ij*_ matrix) in our parameters. In fact, we have the following

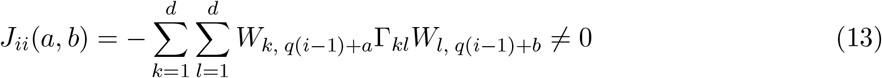

whenever both the patterns (*i, a*) and (*i, b*) are weighted by some principal components that contribute to the energy in the low-dimensional space (Γ_*kl*_ ≠ 0).

### 4.4 Dynamics in the low-dimensional space

Now that we have defined how sequences are projected in the latent space, we need to specify the dynamics that the sequences follow during the laboratory evolution experiment. We choose to model the latent space dynamics as an OU process [33, 34]. The time evolution of the propagator of the trajectory of each sequence from an initial state ***x***_0_ to a sequence **x** at time *t*, is given by the following Fokker-Planck equation:

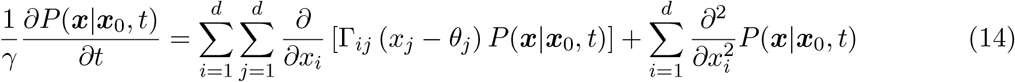

Without loss of generality, we can set the diffusion constant to 1 as it can be absorbed in *γ* and the elements of Γ. The parameter *γ* is the inverse typical time scale of the process, the matrix Γ ∈ ℝ^*d×d*^ defines the external potential that generates the DE trajectory, and the parameter ***θ*** is the stationary point of the dynamics.

Eq. 14 has the following solution:

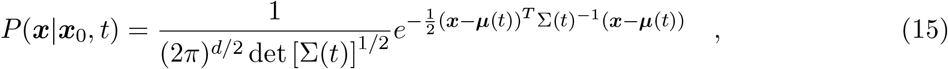

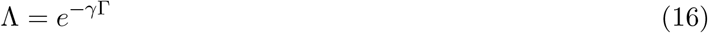

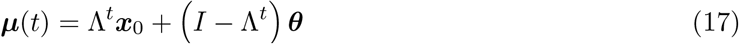

where the time-dependent mean and covariance are parametrized as

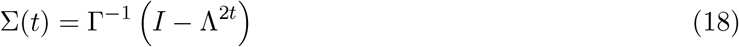

Taking the *t* → ∞ limit, we obtain the equilibrium distribution of our system. It is worth pointing out that the equilibrium solution is still Gaussian and we recover the energy expression from our working hypothesis in Eq. (8).

### 4.5 Maximum Likelihood Inference

Given the latent space dynamics in Eq. (15), we can write the log-likelihood for our data (in our case, the sequencing reads at each evolutionary round). Data are represented as an *M* × *T* matrix and its entries *w*_*m*_(*t*) are the normalized counts of sequence *m* at round *t*. Since the actual pool of sequences appearing in each of the *T* rounds depends on *t*, we indicate with *M* the number all the experimental variant tested in all rounds. Therefore if, for instance, sequence *µ* is not observed at round *t*, then we set *w*_*m*_(*t*) = 0. Let ***a***^*m*^ be the amino acid sequence of variant *m*, ***v***^*m*^ be its one-hot encoded representation, and ***x***^*m*^ = *W* ***v***^*m*^ be its projection onto the latent space. The expression of the (log)-likelihood follows naturally from the model defined in Sec. 4.4.

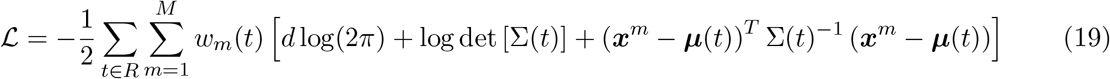

where *R* ⊆ {1, …, *T*} is the set of experimental rounds in which sequencing has been performed.

The quantity in Eq. (19) is numerically maximized as a function of the model parameters {*γ* ∈ ℝ, ***θ*** ∈ ℝ^*d*^, Γ ∈ ℝ^*d×d*^} using the LBFGS algorithm [35–38]. An *l*_2_-regularization, with intensity 0.01, has been added to mitigate over-fitting and avoid parameters becoming too large.

### 4.6 Contact prediction

How to use the Potts model parameters to predict contacts in the three-dimensional structure of the protein is well-established within the DCA approach. It is common practice to score all the possible pairs of sites computing the Frobenius norm of the coupling matrix. For each pair (*i, j*) we evaluate the following quantity:

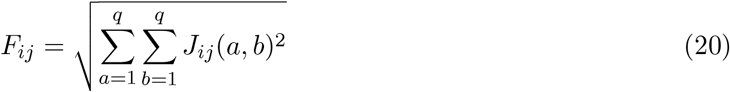

The fact that self-contacts are not considered implies that the couplings in Eq. (13) don’t enter in this score since only *j* ≠ *i* are encountered. This can be a possible reason why one might not worry about this issue.

The quantity in Eq. (20) depends on the gauge of the parameters {*J*_*ij*_(*a, b*)}, we choose the *zero-sum gauge* since it is the one that maximizes this quantity. The zero-sum gauge is the one such that 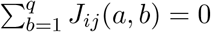 holds for any *i, j* and *a*.

### 4.7 Trajectory reconstruction analysis

To test whether the inferred OU process captures the temporal structure of the directed-evolution experiment, we performed a trajectory reconstruction analysis. This analysis uses the finite-time OU transition density in latent space, defined by Eq. (15), to score experimentally observed variants and compare them with random variants generated from the wild-type sequence. For each latent-space dimension *d*, sequences were first mapped to the PCA latent space as described in Sec. 4.3.

We considered two versions of the analysis. In the first case, the model was trained using the full experimental time series. This setting provides an in-sample consistency check, testing whether the fitted OU dynamics is compatible with the observed evolutionary trajectory. In the second case, the final experimental round was removed from the training set and used only for evaluation. This leave-last-round-out setting provides an out-of-sample test of whether the dynamics inferred from the earlier rounds can recognize experimentally evolved variants.

Let *t*_*∗*_ denote the time of the final experimental round. To reconstruct trajectory from the wild type, each final-round sequence ***x***^*m*^ was assigned the log-transition score divided by *d*

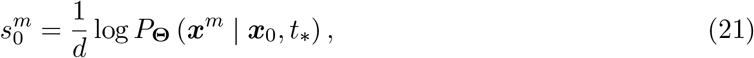

where *P*_**Θ**_ is the OU transition density evaluated with the parameters **Θ** inferred in the corresponding training setting. Here, ***x***_*m*_ = *W* ***v***^*m*^ denotes the latent space representation of sequence *m* from the data; since the variants observed along evolutionary trajectories can be labeled both in the case of experimental data and random sequences, *m* can be though as an integer that indicates a specific sequence from the data collected.

We also computed trajectory reconstruction scores conditioned on earlier experimental rounds using the same expression of the transition probability according to OU dynamics

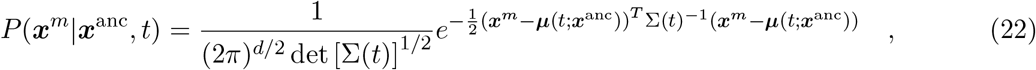

where ***x***^anc^ indicates a generic ancestor sequence that lies along the evolutionary trajectory ending in ***x***^*m*^ at time *t*_*∗*_. Here we explicitely indicated the dependency on ***x***^anc^ of the quantity ***µ***(*t*) defined by Eq. (17). Since individual lineages are not observed in the sequencing data, each final-round sequence was assigned the best transition score over all variants observed at a previous round *t < t*_*∗*_:

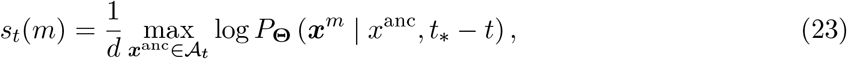

where A_*t*_ is the set of variants observed at round *t*.

This score asks whether a final-round sequence can be dynamically connected to at least one previously observed variant under the learned OU process. As a null model, we generated random sequences from the wild-type sequence using a uniform mutation process. The mutation rate of the random ensemble was matched to the corresponding directed-evolution experiment.

Random and experimental sequences were projected into the same PCA latent space, and the same trajectory reconstruction scores in Eqs. (21) and (23) were computed for both sets. For each value of *d* and each conditioning time *t*, we compared the score distribution of experimental final-round variants

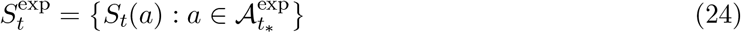

with the score distribution of random variants

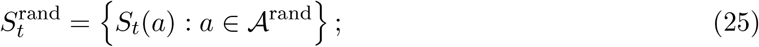

If the model is able to correctly assign log transition probability score, we expect that the values in *S*_exp_ are consistently larger that *S*_rand_ and, consequently, the histogram of the former exhibits a shift towards larger numbers.

Statistical significance was assessed using a one-sided Mann–Whitney *U* test [39], implemented as `MannWhitneyUTest in HypothesisTests.jl` [40]. In our case, the alternative hypothesis is

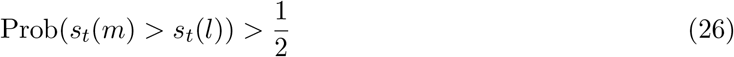

where *m* is a sample uniformly random drawn from the experimental data, and *l* is uniformly random drawn from sequences generated according to the null-model.

Thus, small *p*-values indicate that the experimental final-round variants are assigned higher dynamical likelihood than random variants, while *p*-values close to one indicate that random variants are assigned higher trajectory scores. The resulting *p*-values were analyzed as a function of the latent-space dimension *d* and of the conditioning round *t*. The leave-last-round-out setting was used as the primary validation of predictive trajectory reconstruction, whereas the full-time-series setting was used as an in-sample control.

**Figure 7.**
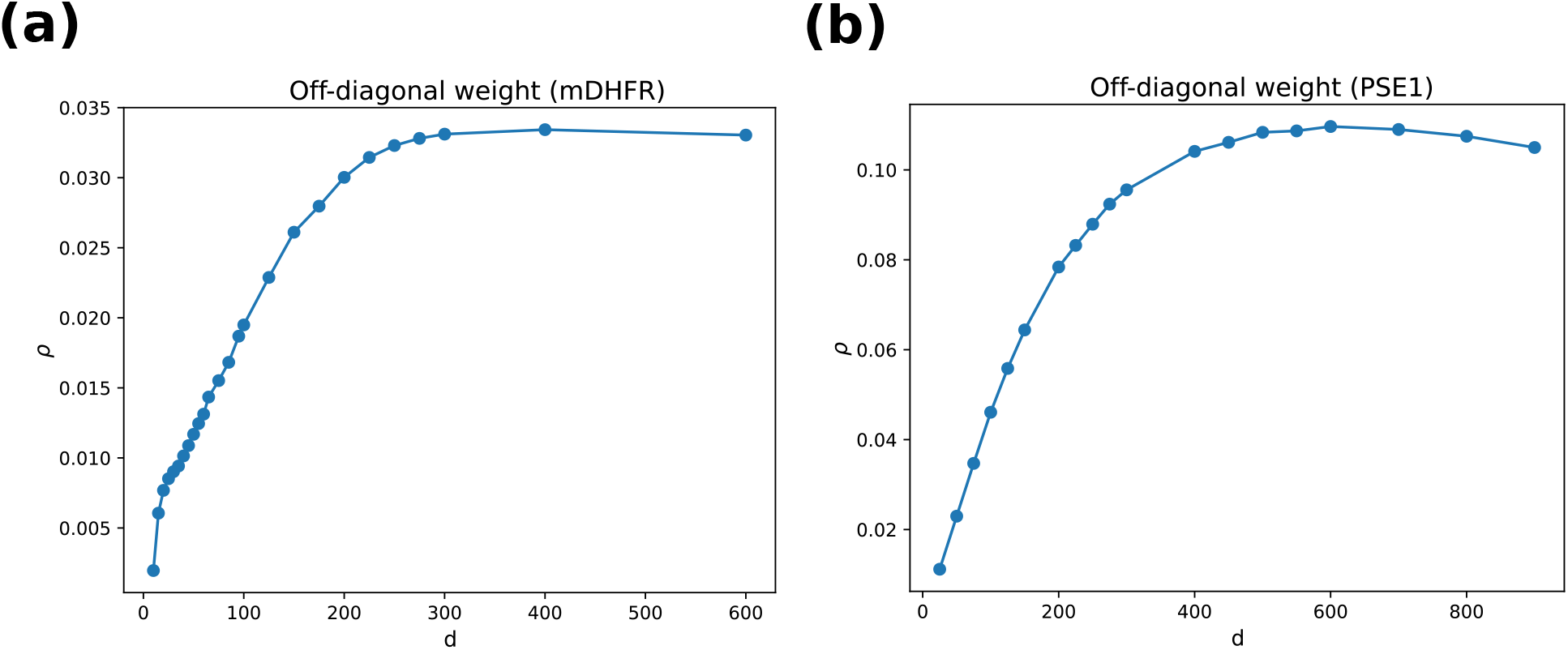
The quantity *ρ* defined by eq. 6 is computed from the model’s parameters for each value of *d*. In both experiments, this quantity exhibits the same behavior: for small *d* it rapidly increases until it reaches saturation.

### 4.8 Latent space dimension

As discussed in sec. 2, it is important to choose a good value for the latent space dimension *d* to obtain significant contact prediction and enrich information from the analysis of natural sequences. Unfortunately, there is no principled and intuitive way to set *d a-priori* or to estimate it from the data, and this is surely the main drawback of our method. Anyway, we have observed that once we have trained our model for several values of *d* we can have an idea if increasing *d* can improve the predictions. We mentioned that one can look at the quantity *ρ* as defined in eq. (6). This quantity intuitively tells us how much co-variation we are adding between PCA patterns that is not present in the natural sequences.

We have mentioned that the trend of this quantity as *d* varies, follows quite accurately the amount of additional contacts, especially long-range ones. This seems to be evident in the case of PSE1 experiment with a strong statistical signal (fig.7 **(b)**). Instead in the case of mDHFR, the picture is more complex and from fig. 7 **(a)** one can see that for very small *d* the quantity *ρ* increases sharply, then at *d* = 75 is continuing to increase but with a smaller slope, and eventually it saturates around *d* = 500. From fig. 2 **(c)** one can notice that around *d* = 75 the OU model predict the maximum amount of additional contacts. This analysis shows that the quantity *ρ* can provide useful information about how to set *d* when the real contact map is not available and we can rely on assessing the model predictions to decide if it is worth to increase *d* or not.

### 4.9 Mapping a discrete set into a continuous space

Rigorously speaking, the dynamical model introduced in Sec. 4.4 is inconsistent with the fact that in the latent space only a finite set of points correspond to valid sequences mapped from the sequence space of one-hot encoded vectors. Eq. 15 instead, assigns a non zero probability to the entire latent space R^*d*^. In our approach, this should be interpreted as a simplifying assumption that allows us to analytically derive the solution for the dynamics as well as the equilibrium distribution of the process. Anyway, a supporting argument for this assumption derived heuristically in the limit of long sequences (*L* → ∞) using the Central Limit Theorem (CLT).

Starting from the equation defining the mapping to the latent space

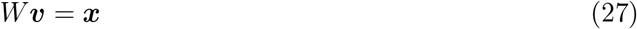

and let us rewrite this equation by splitting the rows of *W* into *L* blocks of size *q. W* is now obtained as horizontal concatenation of *L* matrices *m*^*l*^ of size *d* × *q*, with *l* = {1, …, *L*}.

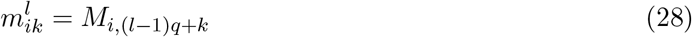

and Eq. 27 becomes

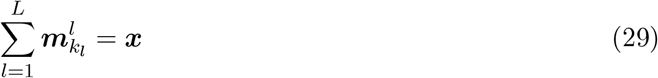

where 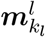 is the vector formed by column *k*_*l*_ of the matrix *m*^*l*^ and *k*_*l*_ are those indices where the one-hot vector ***y*** has entry 1 inside the *l*-th *q*-block.

Let us consider indices *k*_*l*_ as random variables that take values in {1, …, *q*} for each block *l* (for simplicity we assume that they are drawn from an uniform distribution). Then, the following quantity

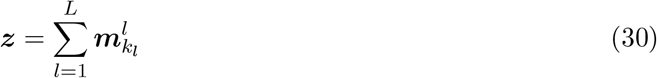

is expressed as the sum of contributions of random variables and, according to the CLT, the following holds in the limit *L* → ∞

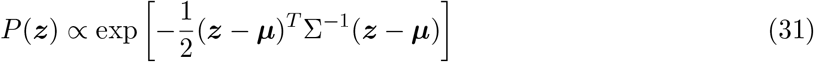

provided the dependence among blocks *m*^*l*^ decays fast enough such that the conditions for the CLT for strong mixing variables hold [41, 42]. The exact expressions of ***µ*** and Σ can be easily derived following the standard computation in the Fourier space that proves the CLT. Eq. (31) states that, in the limit *L* → ∞, *P* (***z***) becomes approximately Gaussian and effectively continuous over the region of the latent space explored by the dynamics. Most of the probability concentrates around ***µ*** and spans over a region of size 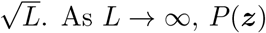 can be approximated locally by a uniform distribution with regions that are small compared to 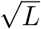.

It is interesting to inspect the relation between the equilibrium distribution of the OU process in the latent space and the equilibrium distribution in the sequence space described by the Potts model. To do this, let us start by writing down the expression of the partition function in the sequence space and use the assumption that the energy of the sequences depends only on their latent representation to do a coarse-graining procedure.

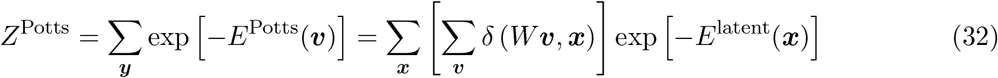

Instead of summing on all possible sequence, we rewrite the expression as a sum over all images in the sequence space. This steps introduces and additional entropic term in the effective energy on the equilibrium distribution defined over the latent space that preserves the Boltzmann weight

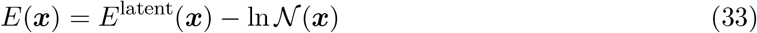

where the additional term ln N(***x***) is defined by

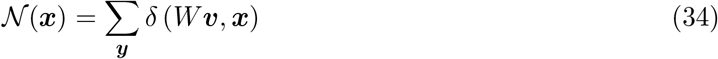

According to the previous argument based on CLT the quantity in Eq. (34) is given by

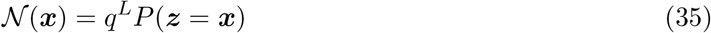

We argued that, in the limit of long sequences, *P* (***z*** = ***x***) can be locally approximated as independent of ***x*** therefore, in the same regime, *N*(***x***) can be considered as approximately constant within the region of the latent space explored by the dynamics. This means that *E*(***x***) and *E*^latent^(***x***) define the same Boltzmann distribution because they differ only by a constant quantity. Therefore, as long as this argument can reasonably be considered valid, the coupling inferred from the equilibrium of the OU process by Eq.3 coincides with the Potts pairwise couplings that describe the interactions among residues in the protein sequence.

## Acknowledgment

MDL, acknowledges interesting discussion with Francesco Zamponi e Saverio Rossi. The Authors acknowledge financial support from the project “Explainable Models for Protein Design”, funded by the MIUR Progetti di Ricerca di Rilevante Interesse Nazionale (PRIN) Bando 2022 - grant 2022TE5B7X. We also acknowledge “Centro Nazionale di Ricerca in High-Performance Computing, Big Data, and Quantum Computing” (ICSC), and “FAIR - Future Artificial Intelligence Research”, received funding from the European Union Next-GenerationEU (Piano Nazionale di Ripresa e Resilienza-Missione 4 Componente 2, Investimento 1.3-D.D. 1555 11/10/2022, PE00000013). This manuscript reflects only the authors’ views and opinions, neither the European Union nor the European Commission can be considered responsible for them.

## References

[1] Adam J. Riesselman, John B. Ingraham, and Debora S. Marks. Deep generative models of genetic variation capture the effects of mutations. Nature Methods, 15(10):816–822, 2018.

[2] Douglas M Fowler and Stanley Fields. Deep mutational scanning: a new style of protein science. Nature Methods, 11(8):801–807, 2014.

[3] Lara Sellés Vidal, Mark Isalan, John T. Heap, and Rodrigo Ledesma-Amaro. A primer to directed evolution: current methodologies and future directions. RSC Chemical Biology, 4(4):271–291, 2023.

[4] Patrick C. Cirino, Kimberly M. Mayer, and Daisuke Umeno. Generating mutant libraries using error-prone pcr. In Directed Evolution Library Creation, pages 3–9. Springer, 2003.

[5] Jesse D. Bloom et al. Protein stability promotes evolvability. Proceedings of the National Academy of Sciences, 103(15):5869–5874, 2006.

[6] Olivier Rivoire. Parsimonious evolutionary scenario for the origin of allostery and coevolution patterns in proteins. Physical Review E, 100(3):032411, 2019.

[7] Jia Zheng, Ning Guo, and Andreas Wagner. Selection enhances protein evolvability by increasing mutational robustness and foldability. Science, 370(6521), 2020.

[8] Kevin K. Yang, Zachary Wu, and Frances H. Arnold. Machine-learning-guided directed evolution for protein engineering. Nature Methods, page 1, 2019.

[9] Philip A. Romero and Frances H. Arnold. Exploring protein fitness landscapes by directed evolution. Nature Reviews Molecular Cell Biology, 10(12):866, 2009.

[10] Marco Fantini, Simonetta Lisi, Paolo De Los Rios, Antonino Cattaneo, and Annalisa Pastore. Protein structural information and evolutionary landscape by in vitro evolution. Molecular Biology and Evolution, 37(4):1179–1192, October 2019.

[11] Michael A. Stiffler, Frank J. Poelwijk, Kelly P. Brock, Richard R. Stein, Adam Riesselman, Joan Teyra, Sachdev S. Sidhu, Debora S. Marks, Nicholas P. Gauthier, and Chris Sander. Protein structure from experimental evolution. Cell Systems, 10(1):15–24.e5, January 2020.

[12] Inferring protein fitness landscapes from laboratory evolution experiments. PLOS Computational Biology, 19(3):e1010956, March 2023.

[13] Luca Sesta, Guido Uguzzoni, Jorge Fernandez-de Cossio-Diaz, and Andrea Pagnani. Amala: Analysis of directed evolution experiments via annealed mutational approximated landscape. International Journal of Molecular Sciences, 22(20):10908, October 2021.

[14] Luca Sesta, Andrea Pagnani, Jorge Fernandez-de Cossio-Diaz, and Guido Uguzzoni. Inference of annealed protein fitness landscapes with annealdca. PLOS Computational Biology, 20(2):e1011812, February 2024.

[15] Muhammad Saqib Sohail, Raymond H. Y. Louie, Matthew R. McKay, and John P. Barton. Mpl resolves genetic linkage in fitness inference from complex evolutionary histories. Nature Biotechnology, 39(4):472–479, Apr 2021.

[16] Muhammad Saqib Sohail, Raymond H Y Louie, Zhenchen Hong, John P Barton, and Matthew R McKay. Inferring Epistasis from Genetic Time-series Data. Molecular Biology and Evolution, 39(10):msac199. 09 2022.

[17] Thomas F. Hansen. Stabilizing selection and the comparative analysis of adaptation. Evolution, 51(5):1341–1351, October 1997.

[18] Russell Lande. Natural selection and random genetic drift in phenotypic evolution. Evolution, 30(2):314–334, 1976.

[19] Edwin Rodríguez Horta, Alejandro Lage-Castellanos, Martin Weigt, and Pierre Barrat-Charlaix. Global multivariate model learning from hierarchically correlated data. Journal of Statistical Mechanics: Theory and Experiment, 2021(7):073501, jul 2021.

[20] Najeeb Halabi, Olivier Rivoire, Stanislas Leibler, and Rama Ranganathan. Protein sectors: Evolutionary units of three-dimensional structure. Cell, 138(4):774–786, August 2009.

[21] Christopher J R Illingworth and Ville Mustonen. Distinguishing driver and passenger mutations in an evolutionary history categorized by interference. Genetics, 189(3):989–1000, November 2011.

[22] Martin Weigt, Robert A. White, Hendrik Szurmant, James A. Hoch, and Terence Hwa. Identification of direct residue contacts in protein–protein interaction by message passing. Proceedings of the National Academy of Sciences, 106(1):67–72, 2009.

[23] Faruck Morcos, Andrea Pagnani, Bryan Lunt, Arianna Bertolino, Debora S. Marks, Chris Sander, Riccardo Zecchina, José N. Onuchic, Terence Hwa, and Martin Weigt. Direct-coupling analysis of residue coevolution captures native contacts across many protein families. Proceedings of the National Academy of Sciences, 108(49), November 2011.

[24] Simona Cocco, Christoph Feinauer, Matteo Figliuzzi, Rémi Monasson, and Martin Weigt. Inverse statistical physics of protein sequences: a key issues review. Reports on Progress in Physics, 81(3):032601, jan 2018.

[25] Jakub Otwinowski, David M. McCandlish, and Joshua B. Plotkin. Inferring the shape of global epistasis. Proceedings of the National Academy of Sciences, 115(32), 2018.

[26] Magnus Ekeberg, Tuomo Hartonen, and Erik Aurell. Fast pseudolikelihood maximization for direct-coupling analysis of protein structure from many homologous amino-acid sequences. Journal of Computational Physics, 276:341–356, November 2014.

[27] Carlo Baldassi, Marco Zamparo, Christoph Feinauer, Andrea Procaccini, Riccardo Zecchina, Martin Weigt, and Andrea Pagnani. Fast and accurate multivariate gaussian modeling of protein families: predicting residue contacts and protein-interaction partners. PloS one, 9(3):e92721, 2014.

[28] Jerome Friedman, Trevor Hastie, and Robert Tibshirani. Sparse inverse covariance estimation with the graphical lasso. Biostatistics, 9(3):432–441, December 2007.

[29] Thomas A Hopf, Anna G Green, Benjamin Schubert, Sophia Mersmann, Charlotta P I Schärfe, John B Ingraham, Agnes Toth-Petroczy, Kelly Brock, Adam J Riesselman, Perry Palmedo, Chan Kang, Robert Sheridan, Eli J Draizen, Christian Dallago, Chris Sander, and Debora S Marks. The evcouplings python framework for coevolutionary sequence analysis. Bioinformatics, 35(9):1582–1584, October 2018.

[30] Sean R. Eddy. Accelerated profile hmm searches. PLoS Computational Biology, 7(10):e1002195. 2011.

[31] Simona Cocco, Remi Monasson, and Martin Weigt. From principal component to direct coupling analysis of coevolution in proteins: Low-eigenvalue modes are needed for structure prediction. PLoS Computational Biology, 9(8):e1003176, August 2013.

[32] Thomas A Hopf, John B Ingraham, Frank J Poelwijk, Charlotta P I Schärfe, Michael Springer, Chris Sander, and Debora S Marks. Mutation effects predicted from sequence co-variation. Nature Biotechnology, 35(2):128–135, January 2017.

[33] Crispin W. Gardiner. Stochastic Methods: A Handbook for the Natural and Social Sciences. Springer, Berlin, Heidelberg, 4 edition, 2009.

[34] G. E. Uhlenbeck and L. S. Ornstein. On the theory of the brownian motion. Physical Review, 36(5):823–841, 1930.

[35] C. G. Broyden. The convergence of a class of double-rank minimization algorithms: 2. the new algorithm. IMA Journal of Applied Mathematics, 6(3):222–231, 1970.

[36] R. Fletcher and M. J. D. Powell. A rapidly convergent descent method for minimization. The Computer Journal, 6(2):163–168, August 1963.

[37] Donald Goldfarb. A family of variable-metric methods derived by variational means. Mathematics of Computation, 24(109):23–26, 1970.

[38] D. F. Shanno. Conditioning of quasi-newton methods for function minimization. Mathematics of Computation, 24(111):647–656, 1970.

[39] Henry B. Mann and Donald R. Whitney. On a test of whether one of two random variables is stochastically larger than the other. The Annals of Mathematical Statistics, 18(1):50–60, 1947.

[40] JuliaStats. Hypothesistests.jl: Hypothesis tests for julia, 2026. Julia package, MannWhitneyUTest.

[41] Walter Kramer. Probability & measure : Patrick billingsley (1995): (3rd ed.). new york : Wiley, isbn 0-471-0071-02, pp 593, [pound sign] 49.95. Computational Statistics & Data Analysis, 20(6):702–703, December 1995.

[42] Richard C. Bradley. Basic properties of strong mixing conditions. a survey and some open questions. Probability Surveys, 2(none), January 2005.

