## Supplementary Information for "Modeling Protein Sequence Evolution as an Ornstein-Uhlenbeck Process in a Latent Space"

### S1 PlmDCA predictions

Figure S1 panels **(a)** and **(b)**, shows the positive predictive value (PPV) as a function of the number of pairs of residues ranked in reverse PlmDCA contact score order, for both mDHFR and PSE1. PlmDCA reaches an excellent prediction accuracy on both datasets. In particular, correct pairs of long-range distances (on the primary sequence) are detected, as shown in panels **(c)** and **(d)**.

### S2 Combined-MSA predictions

The Combined-MSA baseline was obtained by appending experimental variants to the natural multiple sequence alignment and applying PlmDCA with standard sequence reweighting, as intriduced in Sec. 2.1. Since experimental variants derive from a common wild type and are highly similar to one another, they receive low effective weights. As a consequence, the inferred couplings remain dominated by the natural homologs.

Figure S2 shows the correct contact predictions introduced by Combined-MSA that are absent from PlmDCA trained only on natural sequences. The number of additional contacts is limited for both proteins, and particularly small for DHFR. This confirms that simply pooling natural and experimental sequences makes only modest use of the temporal experimental information, motivating the explicit dynamical treatment introduced in the main text.

### S3 Evolutionary Dynamics Predictions

This section provides the complete results of the trajectory-reconstruction analysis introduced in Sec. 2.3 of the main text. We evaluate whether the finite-time

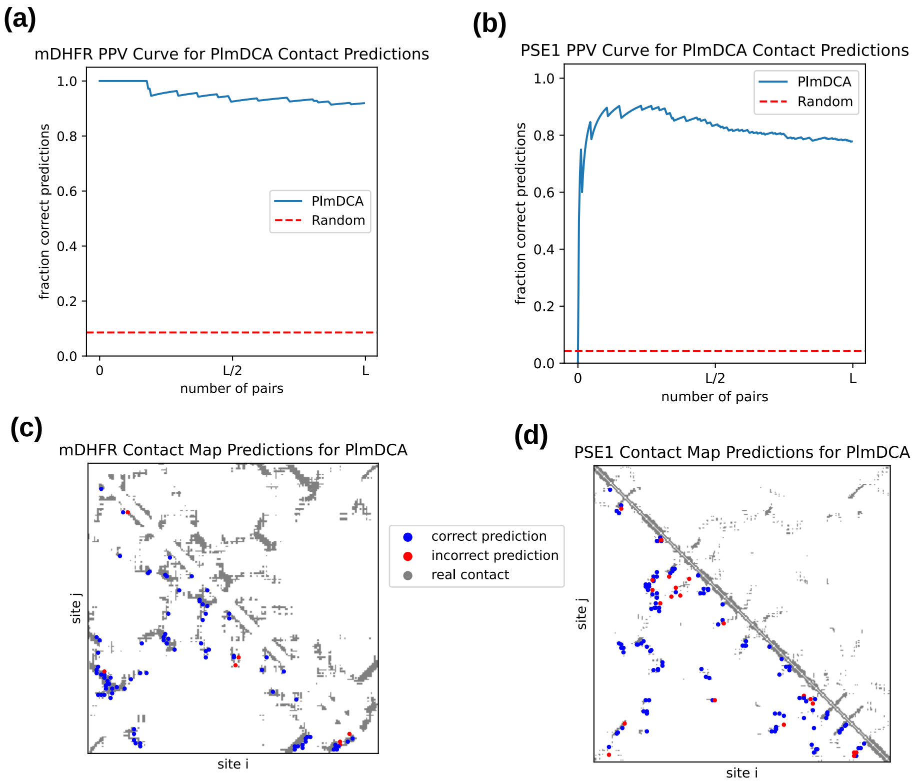


Figure S1: Panels **(a,b)**: Positive Predictive Value (PPV) curves for contact prediction obtained with PlmDCA trained on mDHFR and PSE1 PFAM alignments. The x-axis reports the number of top-ranked site pairs (ordered by decreasing score), while the y-axis shows the fraction of these pairs that correspond to true contacts in the 3D structure.
Panels **(c,d)**: Contact maps for PlmDCA trained on natural sequences of mDHFR and PSE1. The top $L/2$ ranked site pairs are shown. Blue dots indicate correctly predicted contacts (i.e., pairs in contact in the 3D structure), while red dots denote incorrect predictions. The fraction of correct predictions corresponds to the PPV value at $L/2$ in panels **(a,b)**

.

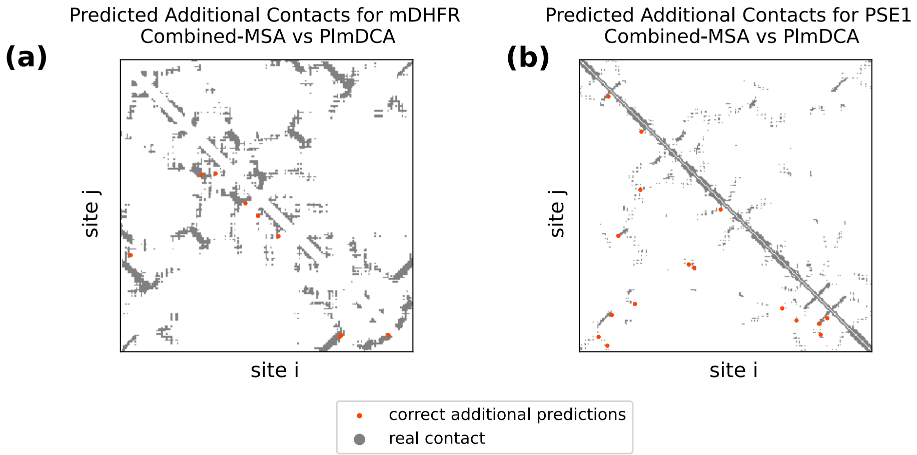


Figure S2: **(a)**: Contact map of the predictions from Combined-MSA that were not found by PlmDCA for DHFR. **(b)**: Same as **(a)** for PSE1.

Ornstein–Uhlenbeck transition kernel assigns higher scores to experimentally observed final-round variants than to variants generated from randomized sequence ensembles.

Two training settings are considered. In the full-time-series analysis, all experimental rounds, including the final round, are used for parameter inference. This analysis is therefore an in-sample consistency check. In the leave-last-round-out analysis, the final experimental round is excluded from model training and used only for evaluation, providing an out-of-sample assessment of the inferred dynamics.

For each latent-space dimension $d$, trajectory scores are computed either directly from the wild-type sequence, corresponding to $t = 0$, or conditionally on variants observed at an earlier experimental round. Since individual genealogical relationships are unavailable, conditional reconstruction uses the maximum transition score over all candidate ancestor sequences observed at the conditioning round. Statistical significance is assessed using a one-sided Mann–Whitney U test, with the alternative hypothesis that experimental variants receive higher scores than randomized controls.

#### S3.1 Uniformly Random Null

In the uniform-mutation null, random variants are generated from the wild-type sequence using the average mutation rate measured in each experimental round. Mutation positions and amino-acid identities are otherwise sampled uniformly. This ensemble approximately preserves the mutational distance from the wild type while removing site-specific preferences and correlations between mutations.

For PSE1, experimental final-round variants are strongly favored over uni-

| $d$ | $t = 0$ | $t = 10$ |
|---|---|---|
| 25 | 0 | 2.31e-31 |
| 50 | 0 | 1.29e-32 |
| 75 | 0 | 1.63e-30 |
| 100 | 0 | 3.38e-30 |
| 125 | 0 | 1.22e-29 |
| 150 | 0 | 5.71e-26 |
| 200 | 0 | 1.09e-28 |
| 225 | 0 | 4.97e-28 |
| 250 | 0 | 1.11e-26 |
| 275 | 0 | 4.47e-28 |
| 300 | 0 | 1.30e-27 |
| 400 | 0 | 2.12e-31 |
| 450 | 0 | 1.15e-29 |
| 500 | 0 | 8.10e-29 |
| 550 | 0 | 2.86e-29 |
| 600 | 0 | 5.09e-29 |
| 700 | 0 | 4.20e-29 |
| 800 | 0 | 3.38e-30 |
| 900 | 0 | 5.91e-27 |

Table S1: Trajectory-reconstruction p-values for **PSE1**, **full time-series training**. The alternative hypothesis is that experimental final-round variants have higher OU transition scores than **uniformly generated random** variants.

formly generated controls both when the model is trained on the full time series and when the final round is excluded from training. In the leave-last-round-out setting, the direct transition from the wild type and the transition conditioned on round 10 remain significant across all tested latent dimensions, indicating that the reconstructed trajectory is not explained by mutation load alone.

For DHFR, the full-time-series analysis also produces strong significance over a broad range of dimensions. However, this result primarily reflects in-sample consistency because the final round contributes to model fitting. In the held-out analysis, significant reconstruction is restricted to a narrow set of dimensions, particularly around $d = 125$ and $d = 150$. At most other dimensions, the one-sided p-values are close to one, showing that the model does not robustly generalize to the held-out DHFR population.

The contrast between the two datasets is consistent with their different experimental diversity. The broader mutational exploration observed in PSE1 provides a stronger temporal signal, whereas the limited variation between DHFR rounds makes the inferred dynamics more sensitive to the selected PCA representation.

| $d$ | $t = 0$ | $t = 10$ |
|---|---|---|
| 10 | 0 | 9.82e-31 |
| 25 | 0 | 4.91e-32 |
| 50 | 0 | 2.16e-28 |
| 75 | 0 | 1.20e-27 |
| 100 | 0 | 1.30e-26 |
| 125 | 0 | 1.30e-27 |
| 150 | 0 | 5.98e-24 |
| 175 | 0 | 1.06e-25 |
| 200 | 0 | 2.03e-22 |
| 225 | 0 | 2.05e-21 |
| 250 | 0 | 1.33e-25 |
| 275 | 0 | 5.81e-23 |
| 300 | 0 | 4.12e-24 |

Table S2: Trajectory-reconstruction p-values for **PSE1**, **leave-last-round-out**. The alternative hypothesis is that experimental final-round variants have higher OU transition scores than **uniformly generated random** variants.

#### S3.2 Site-Dependent Mutation Rate Null

The uniformly random null does not account for the fact that different sequence positions can have substantially different mutation probabilities. To construct a more stringent control, we estimate for each experimental round the empirical probability $f_i^{\mathrm{mut}}$ that site $i$ differs from the wild-type residue. Random sequences are then generated independently across positions according to $P(a_i \neq a_i^{\mathrm{WT}}) = f_i^{\mathrm{mut}}$, while the identity of the mutant amino acid is sampled uniformly among the amino acids different from the wild-type residue. This null preserves the position-dependent mutation rate but removes amino-acid-specific preferences and correlations between sites.

The site-dependent null substantially reduces the trajectory-reconstruction signal. For PSE1, the held-out experimental variants are not significantly favored across most low- and intermediate-dimensional representations. Significant discrimination appears only for the largest tested latent spaces. For the direct wild-type-to-final-round reconstruction, significant p-values are obtained at $d =$ 400, 500, 600 and 700. When conditioning on round 10, the model fails in recognizing experimental sequences from random control. These results suggest that information beyond site-specific mutation propensity is present in PSE1, but it is distributed across a relatively large number of latent directions. A plausible explanation for this failure is that, when the last experimental round is removed, the model is trained only on two time point snapshots ($t = 0$, and $t = 10$); therefore, all transitions start from the wild-type sequence and there is not enough information to correctly estimate the general dynamics of the process from other initial sequences (see Tab. S5).

In contrast, no clear signal is observed for DHFR under the site-dependent null. This is consistent with the limited temporal variation of the DHFR libraries: once site-specific mutation frequencies are reproduced, little additional information remains for the OU dynamics to distinguish the experimental final round from the randomized ensemble.

Compared with the uniform null, this analysis supports a more restricted conclusion. The strong uniform-null results show that the model captures the nonuniform organization of experimentally explored sequence space. For high-dimensional PSE1 representations, the model captures sequence features that are not reproduced by a null model preserving only the site-specific mutation probabilities.

#### S3.3 Site-Dependent Profile Null

As a still more constrained control, we generated sequences by independently sampling the amino acid at each position from the empirical single-site frequency profile of the corresponding experimental round. This null therefore preserves the complete marginal amino-acid distribution at every site while removing correlations between different positions.

In contrast to the site-dependent mutation-rate null, the profile null reproduces not only the probability that each site is mutated but also the amino-acid preferences among mutant residues. After projection into the PCA latent space, this ensemble closely reproduces the experimental mean and much of the covariance structure. Consequently, experimental and profile-randomized variants are generally not significantly separated by the OU transition score (Fig. S3 reports an example for $d = 600$).

This negative result is informative because it clarifies the origin of the trajectory-reconstruction signal. The model robustly distinguishes experimental variants from controls preserving only the total mutation rate and, for high-dimensional PSE1 representations, from controls preserving position-specific mutation probabilities. However, it does not robustly distinguish them from controls preserving the full set of single-site amino-acid frequencies. Thus, a substantial fraction of the predictive signal is encoded in single-site sequence statistics rather than in higher-order correlations alone.

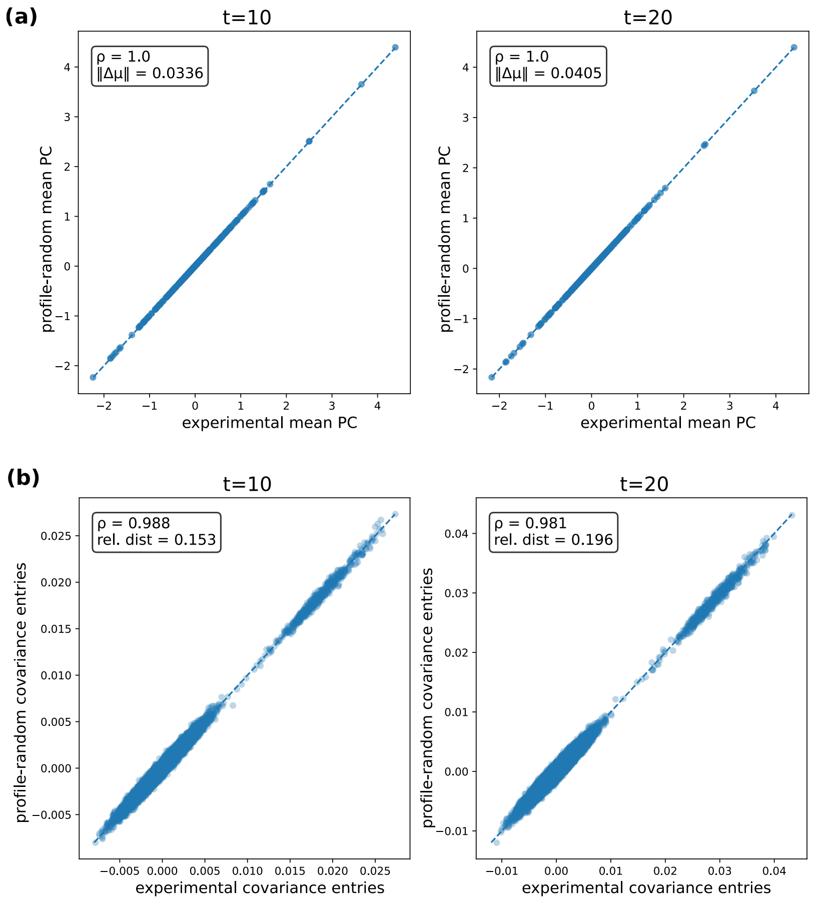


Figure S3: Panel **(a)** reports a comparison between the latent-space coordinates of the mean of the experimental sequences and those obtained from profile-randomized sequences at $t = 10$ and $t = 20$. Each point represents one principal component; the dashed line denotes equality. The Pearson correlation coefficient $\rho$ and the Euclidean norm of the difference between the mean vectors, $||\Delta\mu||$, are reported in each panel. In panel **(b)** an analogous comparison for the entries of the covariance matrices. Each point represents one covariance-matrix element, and the dashed line denotes equality. The panels report the Pearson correlation coefficient and the relative distance between the experimental and profile-randomized covariance matrices. The near-perfect agreement shows that profile randomization preserves both the mean structure and, to a large extent, the covariance structure of the experimental data after projection into the latent PCA space.

| $d$ | $t=0$ | $t=1$ | $t=2$ | $t=3$ | $t=4$ | $t=5$ |
|---|---|---|---|---|---|---|
| 15 | 9.99e-01 | 7.81e-01 | 7.81e-01 | 7.81e-01 | 7.81e-01 | 7.81e-01 |
| 20 | 3.79e-01 | 9.10e-01 | 9.10e-01 | 9.10e-01 | 9.10e-01 | 9.10e-01 |
| 25 | 3.08e-06 | 2.81e-01 | 2.81e-01 | 2.81e-01 | 2.81e-01 | 2.81e-01 |
| 30 | 1.71e-35 | 1.72e-01 | 1.72e-01 | 1.72e-01 | 1.72e-01 | 1.72e-01 |
| 40 | 4.79e-224 | 8.69e-13 | 8.69e-13 | 8.69e-13 | 8.69e-13 | 8.69e-13 |
| 45 | 1.99e-308 | 2.93e-16 | 2.93e-16 | 2.93e-16 | 2.93e-16 | 2.93e-16 |
| 50 | 0 | 1.30e-22 | 1.30e-22 | 1.30e-22 | 1.30e-22 | 1.30e-22 |
| 55 | 0 | 3.36e-25 | 3.36e-25 | 3.36e-25 | 3.36e-25 | 3.36e-25 |
| 60 | 0 | 2.76e-31 | 2.76e-31 | 2.76e-31 | 2.76e-31 | 2.76e-31 |
| 65 | 0 | 2.43e-19 | 2.43e-19 | 2.43e-19 | 2.43e-19 | 2.43e-19 |
| 70 | 0 | 3.29e-22 | 3.29e-22 | 3.29e-22 | 3.29e-22 | 3.29e-22 |
| 75 | 0 | 1.72e-29 | 1.72e-29 | 1.72e-29 | 1.72e-29 | 1.72e-29 |
| 80 | 0 | 3.01e-31 | 3.01e-31 | 3.01e-31 | 3.01e-31 | 3.01e-31 |
| 85 | 0 | 1.77e-29 | 1.77e-29 | 1.77e-29 | 1.77e-29 | 1.77e-29 |
| 90 | 0 | 1.51e-31 | 1.51e-31 | 1.51e-31 | 1.51e-31 | 1.51e-31 |
| 95 | 0 | 1.16e-32 | 1.16e-32 | 1.16e-32 | 1.16e-32 | 1.16e-32 |
| 100 | 0 | 3.61e-39 | 3.61e-39 | 3.61e-39 | 3.61e-39 | 3.61e-39 |
| 125 | 0 | 3.22e-40 | 3.22e-40 | 3.22e-40 | 3.22e-40 | 3.22e-40 |
| 150 | 0 | 1.53e-40 | 1.53e-40 | 1.53e-40 | 1.53e-40 | 1.53e-40 |
| 175 | 0 | 3.34e-49 | 3.34e-49 | 3.34e-49 | 3.34e-49 | 3.34e-49 |
| 200 | 0 | 1.97e-57 | 1.97e-57 | 1.97e-57 | 1.97e-57 | 1.97e-57 |
| 225 | 0 | 1.02e-55 | 1.02e-55 | 1.02e-55 | 1.02e-55 | 1.02e-55 |
| 250 | 0 | 3.66e-62 | 3.66e-62 | 3.66e-62 | 3.66e-62 | 3.66e-62 |
| 275 | 0 | 2.56e-70 | 2.56e-70 | 2.56e-70 | 2.56e-70 | 2.56e-70 |
| 300 | 0 | 2.86e-41 | 2.87e-41 | 2.86e-41 | 2.85e-41 | 2.85e-41 |
| 350 | 0 | 2.08e-78 | 2.08e-78 | 2.08e-78 | 2.08e-78 | 2.08e-78 |
| 400 | 0 | 2.37e-78 | 2.37e-78 | 2.37e-78 | 2.37e-78 | 2.37e-78 |
| 500 | 0 | 1.32e-96 | 1.32e-96 | 1.32e-96 | 1.32e-96 | 1.32e-96 |

Table S3: Trajectory-reconstruction p-values for **DHFR**, **full time-series training**. The alternative hypothesis is that experimental final-round variants have higher OU transition scores than **uniformly generated random** variants.

| $d$ | $t = 0$ | $t = 1$ | $t = 2$ | $t = 3$ | $t = 4$ | $t = 5$ |
|---|---|---|---|---|---|---|
| 10 | 1.00e+00 | 9.99e-01 | 9.99e-01 | 9.99e-01 | 9.99e-01 | 9.99e-01 |
| 25 | 1.00e+00 | 9.98e-01 | 9.98e-01 | 9.98e-01 | 9.98e-01 | 9.98e-01 |
| 50 | 1.00e+00 | 2.27e-01 | 2.27e-01 | 2.27e-01 | 2.27e-01 | 2.27e-01 |
| 75 | 1.00e+00 | 7.29e-01 | 9.28e-01 | 2.71e-01 | 1.37e-01 | 1.75e-02 |
| 100 | 1.00e+00 | 9.38e-01 | 9.38e-01 | 9.38e-01 | 9.38e-01 | 9.38e-01 |
| 125 | 1.39e-55 | 1.35e-05 | 1.35e-05 | 1.35e-05 | 1.35e-05 | 1.35e-05 |
| 150 | 6.63e-28 | 2.16e-02 | 2.16e-02 | 2.16e-02 | 2.16e-02 | 2.16e-02 |
| 175 | 1.00e+00 | 1.00e+00 | 1.00e+00 | 1.00e+00 | 1.00e+00 | 9.96e-01 |
| 200 | 1.00e+00 | 1.00e+00 | 1.00e+00 | 1.00e+00 | 1.00e+00 | 1.00e+00 |
| 225 | 1.00e+00 | 1.00e+00 | 1.00e+00 | 1.00e+00 | 1.00e+00 | 1.00e+00 |
| 250 | 1.00e+00 | 1.00e+00 | 1.00e+00 | 1.00e+00 | 1.00e+00 | 1.00e+00 |
| 275 | 1.00e+00 | 1.00e+00 | 1.00e+00 | 1.00e+00 | 1.00e+00 | 1.00e+00 |
| 300 | 1.00e+00 | 1.00e+00 | 1.00e+00 | 1.00e+00 | 1.00e+00 | 1.00e+00 |

Table S4: Trajectory-reconstruction p-values for **DHFR**, **leave-last-round-out**. The alternative hypothesis is that experimental final-round variants have higher OU transition scores than **uniformly generated random** variants.

| $d$ | $t = 0$ | $t = 10$ |
|---|---|---|
| 10 | 1.00e+00 | 1.00e+00 |
| 25 | 1.00e+00 | 1.00e+00 |
| 50 | 1.00e+00 | 1.00e+00 |
| 75 | 1.00e+00 | 9.62e−01 |
| 100 | 1.00e+00 | 1.00e+00 |
| 125 | 1.00e+00 | 1.00e+00 |
| 150 | 6.10e−01 | 5.63e−01 |
| 175 | 1.00e+00 | 9.80e−01 |
| 200 | 1.00e+00 | 9.30e−01 |
| 225 | 1.00e+00 | 8.35e−02 |
| 250 | 9.99e−01 | 9.67e−01 |
| 275 | 1.00e+00 | 9.29e−01 |
| 300 | 1.00e+00 | 6.65e−01 |
| 400 | 3.24e−28 | 1.66e−01 |
| 500 | 3.73e−41 | 1.20e−01 |
| 600 | 7.64e−33 | 3.77e−01 |
| 700 | 2.48e−25 | 7.17e−01 |

Table S5: Trajectory-reconstruction p-values for **PSE1** in the **leave-last-round-out** setting using the **site-dependent mutation-rate** null model. The alternative hypothesis is that experimental final-round variants have higher OU transition scores than random variants generated while preserving the empirical mutation probability of each site. The column $t = 0$ corresponds to reconstruction directly from the wild type, whereas $t = 10$ corresponds to reconstruction conditioned on variants observed at round 10.
